# Spatial cell disparity in the colonial choanoflagellate *Salpingoeca rosetta*

**DOI:** 10.1101/653519

**Authors:** Benjamin Naumann, Pawel Burkhardt

## Abstract

Choanoflagellates are the closest unicellular relatives of animals (Metazoa). These tiny protists display complex life histories that include sessile as well as different pelagic stages. Some choanoflagellates have the ability to form colonies as well. Up until recently, these colonies have been described to consist of mostly identical cells showing no spatial cell differentiation, which supported the traditional view that spatial cell differentiation, leading to specific cell types in animals, evolved after the split of the last common ancestor of the Choanoflagellata and Metazoa. The recent discovery of single cells in colonies of the choanoflagellate *Salpingoeca rosetta* that exhibit unique cell morphologies challenges this traditional view. We have now reanalyzed TEM serial sections, aiming to determine the degree of similarity of *S. rosetta* cells within a rosette colony. We investigated cell morphologies and nuclear, mitochondrial and food vacuole volumes of 40 individual cells from four different *S. rosetta* rosette colonies and compared our findings to previously published data on sponge choanocytes. Our analysis show that cells in a choanoflagellate colony differ from each other in respect to cell morphology and content ratios of nuclei, mitochondria and food vacuoles. Furthermore, cell disparity within *S. rosetta* colonies is higher compared to cell disparity within sponge choanocytes. Moreover, we discovered the presence of plasma membrane contacts between colonial cells in addition to already described intercellular bridges and filo-/pseudopodial contacts. Our findings indicate that the last common ancestor of Choanoflagellata and Metazoa might have possessed plasma membrane contacts and spatial cell disparity during colonial life history stages.

## Introduction

The development from a fertilized egg cell, the so-called zygote, to an embryo made up by hundreds of cells or to a juvenile and adult consisting of more than thousands to billions of cells is a hallmark of animals (Metazoa). Metazoan development is a complex process that is facilitated by the highly coordinated interplay of several not less complex sub-processes such as cell division (cleavage), cell-cell interaction, cell migration and cell differentiation (Fairclough et al. 2013; Alberts et al. 2014; Brunet and King 2017). The result of this interplay is a multicellular organism consisting of functionally specialized cells, so-called cell types. Diverse cell types are described in non-bilaterian metazoans such as sponges (Porifera), comb jellies (Ctenophora), *Trichoplax* (Placozoa) and jellyfish (Cnidaria) (Sebé-Pedrós, Chomsky, et al. 2018; Sebé-Pedrós, Saudemont, et al. 2018). Many cell types of non-bilaterian metazoans are multifunctional such as pinacocytes in sponges, epithelial muscle cells in cnidarians (protection, contraction), and the “ocellus” in sponge larvae, a single cell that performs locomotor (steering), photoreceptive and pigmentation functions (Wagner 2014). In bilaterian metazoans multifunctional cells are often replaced by more specialized cell types that sometimes exert only one, very specific, function (Wagner 2014).

Collar cells, polarized cells with an apical flagellum surrounded by a microvillar collar (Leadbeater 2015), are multifunctional cell types (Arendt et al. 2016; Brunet and King 2017; Arendt et al. 2019) and conserved across almost all metazoans and their closest relatives, the choanoflagellates (Brunet and King 2017; Laundon et al. 2019) (Figure 1A). The colony-forming choanoflagellate *Salpingoeca rosetta* (Dayel et al., 2011) has emerged as a promising model organism to investigate the evolutionary origin of metazoan multicellularity and cell differentiation (Dayel et al. 2011; Hoffmeyer and Burkhardt 2016; Booth et al. 2018). *S. rosetta* exhibits a complex life history including different single cell and colony morphologies (Dayel et al. 2011) (Figure 1B). Rosette colony formation is induced by RIF (rosette inducing factor), a sulfonolipid secreted by the bacterium *Algoriphagus machipongonensis* (Alegado et al. 2012; Woznica et al. 2016). Similar to metazoans, colonies of *S. rosetta* form by mitotic divisions from a single founder cell. Cells within a rosette colony are held together by intercellular cytoplasmatic bridges, filopodia and an extracellular matrix (Dayel et al 2011; Laundon et al. 2019).

**Figure 1:**
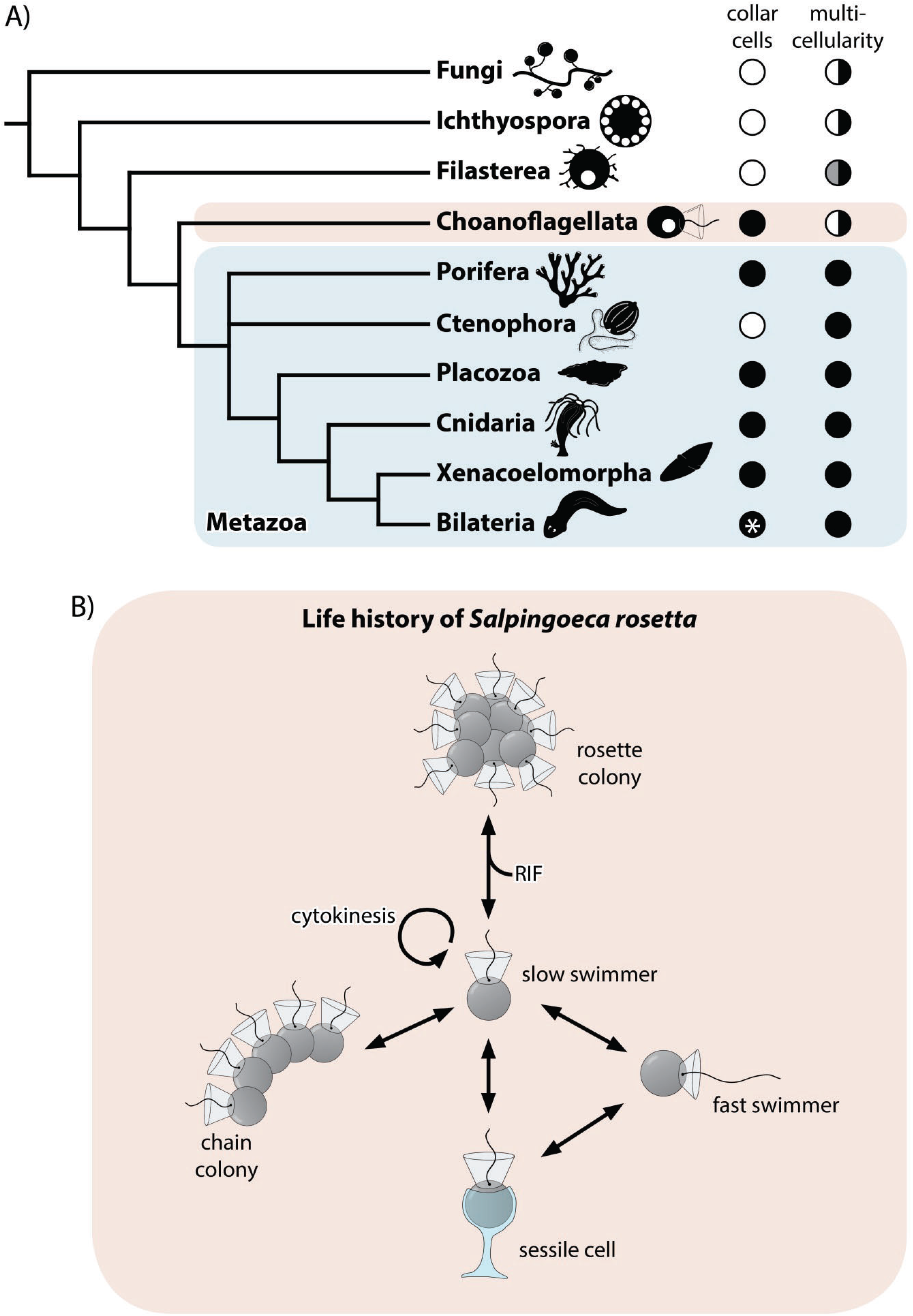
A, Phylogenetic tree showing Choanoflagellata as sister group of the Metazoa (Steenkamp et al. 2005; Carr et al. 2008; Ruiz-Trillo et al. 2008; Adl et al. 2019). In addition, the presence (black circle) and absence (white circle) of collar cells and multicellularity across lineages are shown (adapted from Brunet and King 2017; Laundon et al. 2019). The white asterisk indicates independent secondary losses. Half-filled circles indicate multicellularity in only some species. In Filasterea, multicellularity is achieved by aggregation of single cells (half-filled grey-black circle) instead of clonal division. B, Life history of the choanoflagellate *S. rosetta* after Dayel et al. (2011). RIF, rosette inducing factor (Alegado et al. 2012).

Whether cells of a rosette colony represent a cluster in which cells are identical to each other or colonial differ from each other is still unclear. Although transcriptomic analyses have shown nearly identical expression patterns for single and colonial cells in *S. rosetta*, (Fairclough et al. 2013), a recent study described a distinct morphology of cells in some *S. rosetta* colonies indicating differences of cells within choanoflagellate colonies (Laundon et al. 2019). Understanding whether individual cells of a choanoflagellate colony are identical or different from each other is important for a better understanding of the evolutionary origin of the Metazoa. There are several theories on the evolutionary origin of metazoan multicellular development, cell differentiation and cell types (Fairclough et al. 2010; Sebe-Pedros et al. 2017). As proposed by Ernst Haeckel under the term “Blastaea/Gastraea theory” (Haeckel 1874), animals evolved through “…repeated self-division of [a] primary cell,…” (Haeckel 1892). In this scenario, the last common ancestor of the Metazoa originated from incomplete cell division of a primary single cell that formed a ball-shaped colony called a “Blastaea” consisting of identical cells (Mikhailov et al. 2009). During evolution, intra-colonial division of labor led to increasing differences between cells resulting in specialized cells representing distinct cell types (King 2004). In this scenario, spatial cell differentiation evolved before temporal cell differentiation in the stem lineage of the Metazoa (Mikhailov et al. 2009). A contradicting hypothesis was proposed by Alexey Zakhvatkin in 1949 (Zakhvatkin 1949). The so-called “Synzoospore theory” claimed that metazoans evolved from a unicellular ancestor that showed a variety of different cell types during different life history stages. According to this theory, temporal cell differentiation was already present and accompanied by spatial cell differentiation in the metazoan stem lineage (Mikhailov et al. 2009).

In this study, we used ultrathin transmission electron microscopy (ssTEM) serial sections of whole rosette colonies of *S. rosetta* (previously published by Laundon et al. 2019) to prepare three-dimensional (3D) reconstructions and measure volumes of cell bodies, nuclei, food vacuoles and mitochondria of 40 individual colonial cells from four colonies. We compared the cellular anatomy of colonial *S. rosetta* cells with available data (Laundon et al. 2019) on the cellular anatomy of choanocytes of the homoscleromorph sponge *Oscarella carmela* (Ereskovsky et al. 2017). This comparison allowed us to reveal whether (1) cells in *S. rosetta* rosette colonies are indeed identical (presence of spatial cell disparity), (2) how variable the volume ratios of different cellular organelles are within complete rosette colonies (degree of spatial disparity) and (3) if a similar degree of spatial cell disparity is present in sponge choanocytes (representing a distinct metazoan cell type).

## Materials and Methods

### 3D reconstructions of complete *S. rosetta* colonies

A summary of the workflow is shown in Supplementary Figure 1 (S1). For our analysis, we used digital image stacks of TEM sections of complete *S. rosetta* colonies (RC1 to RC4; n = 40 cells), previously published by Laundon et al. 2019 and available from figshare (doi.org/10.6084/m9.figshare.7346750.v2). The image stacks were imported into AMIRA (FEI Visualization Sciences Group) and aligned manually. Subsequently, the cell body and major cell organelles (nucleus, mitochondria, and food vacuoles) were segmented manually. For surface reconstructions, surface models were rendered from the segmented materials, numbers of polygons were reduced and the surfaces were smoothened for the first time. Materials were then imported into Maya (Autodesk), smoothened twice and colored for final image rendering. For volume renderings, segmented materials were subtracted from the main image stack and exported as separate image stacks. Volume renderings of cells and organelles were prepared using VG Studio Max 2.0 (Volume Graphics).

### Surface measurements and volume calculations

Separated image stacks of cell bodies, nuclei, mitochondria and food vacuoles of the cells of RC1 to RC4 were analyzed with Fiji. Image stacks were imported and masked to create a binary image of the cell body or organelle (black) against a white background. The number of black pixels was counted and the corresponding surface areas were measured on each section. Surface area analyses were conducted using unsmoothed, unprocessed materials. Subsequently, surface area measurements were exported to Microsoft Excel 2010 (Microsoft Corporation) and volumes were calculated by multiplying each surface value with the section thickness (RC1: 70 nm; RC2-RC4: 150 nm) and volume ratio calculations and diagrams were prepared.

## Results

### Cells within rosette colonies of *S. rosetta* exhibit a variety of different morphologies

The individual cells of the four investigated rosette colonies (RC1 – RC4) exhibit a variety of volumes/sizes and morphologies (Figure 2). Three-dimensional reconstructions of all cells of RC1 to RC4 are depicted in Supplementary Figures S2 to S5. Many cells exhibit an ovoid morphology, slightly elongated along the apical-basal axis (AB-axis) (Figure 2A). However, some cells exhibit a more roundish (Figure 2B) or ovoid shape horizontally to the AB-axis (Figure 2C). Laundon et al. (2019) described two cells with a distinct morphology within rosette colonies, C5 of RC3 (carrot-like cell; Figure 2D) and C5 of RC4 (chili-like cell; Figure 2E) and our new reconstructions confirm these findings. In addition, our results confirm the findings of Laundon et al. (2019) regarding the presence of cell membrane protrusions for each cell within four different rosette colonies (n = 40 cells). All cells within a rosette colony exhibit a variety of cell membrane protrusions such as filopodia, pseudopodia and larger lobopodia-like protrusions that might represent pino- and/or endocytotic events (Figure 2F-I).

**Figure 2:**
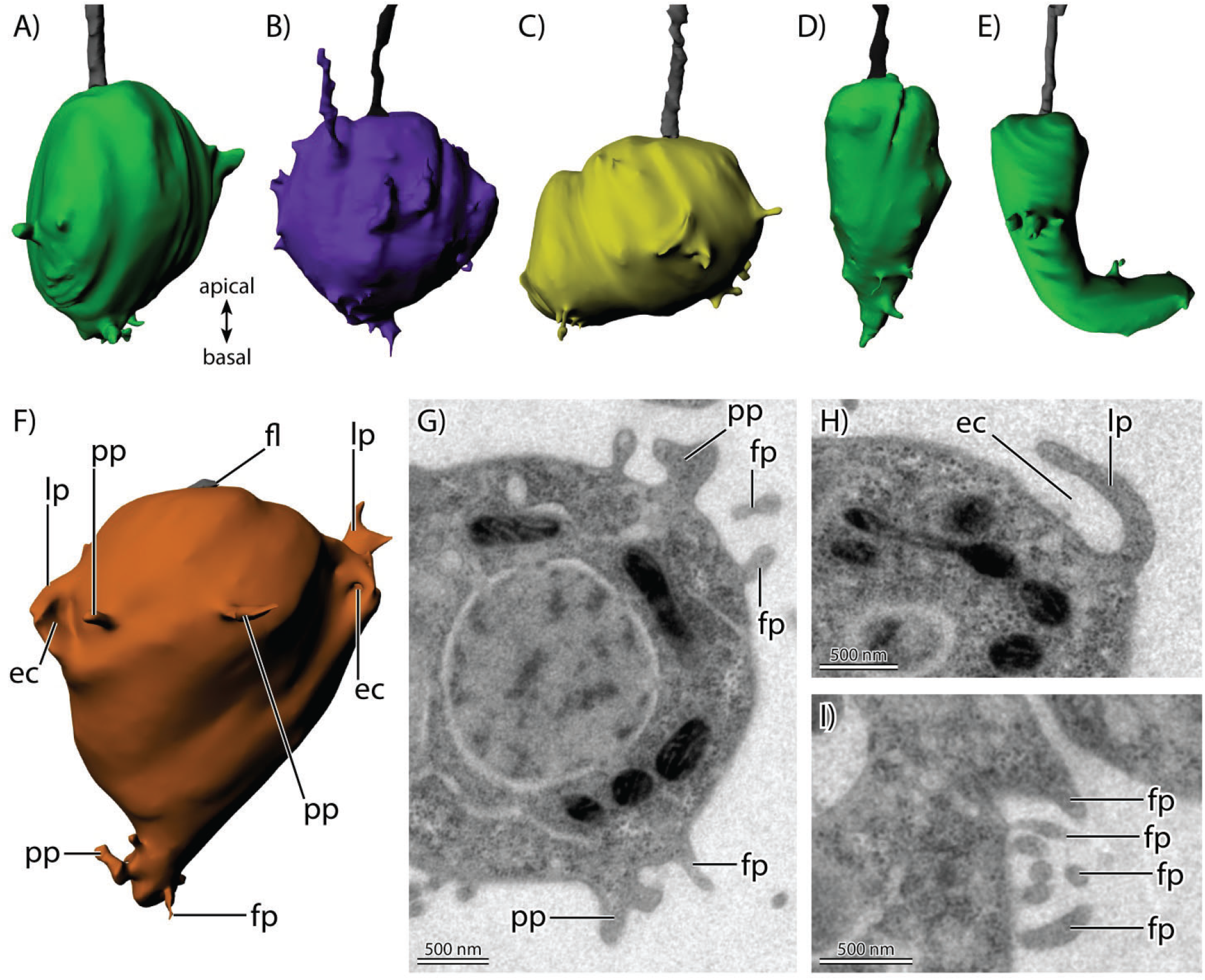
A-F, 3D-surface-reconstructions of cells from four different *S. rosetta* colonies. Cell sizes are not to scale. A, ovoid morphology (RC1, C6). B, roundish morphology (RC3, C10). C, ovoid morphology with the ovoid axis horizontally to the apical-basal axis (RC1, C3). D, “carrot”-cell (RC3, C5). E, “chili”-cell (RC4, C5). F, different types of membrane protrusions within rosette colonies (RC1, C2). G-I, TEM sections of different types of membrane protrusions. ec, endo- or pinocytosis; fl, flagellum; fp, filopodium; lp, lobopodium; pp, pseudopodium.

In RC1 (seven cells; Supplementary Figure S2), cell volumes range from 15,9781 µm^3^ (C2) to 37,711 µm^3^ (C5). Five cells exhibit a more ovoid morphology. Three of these cells (C1, C2, C6) are slightly elongated along the AB-axis. The other two cells (C3, C4) are elongated horizontally to the AB-axis. Two cells (C5, C7) show a more roundish shape

In RC2 (11 cells; Supplementary Figure S3), cell volumes range from 19,1235 µm^3^ (C11) to 46,5795 µm^3^ (C10). Nine cells exhibit a more ovoid morphology. Eight of these cells (C1, C2, C4, C5, C6, C7, C9, C10) are slightly elongated along the AB-axis. C8 is elongated horizontally to the AB-axis. Two cells (C3, C11) show a more roundish shape.

In RC3 (12 cells; Supplementary Figure S4), cell volumes range from 10,1994 µm^3^ (C5) to 51,2403 µm^3^ (C7). Nine cells exhibit a more ovoid morphology. Eight of these cells (C1, C3, C5, C6, C7, C8, C9, C11, C12) are slightly elongated along the AB-axis. As previously reported, C5 exhibits a distinct slender, carrot-like morphology (Laundon et al. 2019). C4 is elongated horizontally to the AB-axis. Two cells (C2, C10) are more roundish. C7, the largest cell of the colony, and C12 show an exceptional high number of membrane protrusions (Supplementary Figure. 4).

In RC4 (10 cells; Supplementary Figure S5), cell volumes range from 13,9838 µm^3^ (C5) to 36,4673 µm^3^ (C8). Seven cells (C1, C3, C4, C6, C7, C9, C10) exhibit a more ovoid morphology, slightly elongated along the AB-axis. As previously reported, C5 exhibits a distinct slender, chili-like morphology (Laundon et al. 2019). Two cells (C2, C8), show a more roundish shape

### Nuclear volume correlates with cell size in *S. rosetta* rosette colonies

In most cells of the four analyzed rosette colonies, the nucleus is located approximately in the middle of the apical-basal axis of the cell. (Figure 3A). For the relative and absolute volume calculations all sub-structures of the nucleus (the nuclear lamina, eu- and heterochromatin and the nucleolus) were included (Table 1).

**Table 1:** Absolute and relative nuclear volumes of cells of RC1 to RC4. abs. V_cell_, absolute total cellular volume; abs. V_nu_, absolute nuclear volume; rel. V_nu_, relative nuclear volume.

| RC1 | C2 | C1 | C6 | C7 | C3 | C4 | C5 |
| --- | --- | --- | --- | --- | --- | --- | --- |
| abs. $V_{\text{cell}}$ in $\mu\text{m}^3$ | 15,9781 | 18,8541 | 22,1971 | 22,5861 | 29,7504 | 36,4994 | 37,7110 |
| abs. $V_{\text{nu}}$ in $\mu\text{m}^3$ | 2,4168 | 3,0355 | 3,2570 | 3,2711 | 4,5026 | 4,8618 | 5,4945 |
| rel. $V_{\text{nu}}$ in % | 15,13 | 16,10 | 14,67 | 14,48 | 15,13 | 13,32 | 14,57 |

| RC2 | C11 | C8 | C2 | C3 | C4 | C7 | C5 | C9 | C6 | C1 | C10 |
| --- | --- | --- | --- | --- | --- | --- | --- | --- | --- | --- | --- |
| abs. $V_{\text{cell}}$ in $\mu\text{m}^3$ | 19,1235 | 21,7361 | 22,5701 | 24,5022 | 24,6890 | 24,7695 | 25,3476 | 27,6794 | 27,7535 | 35,3241 | 46,5795 |
| abs. $V_{\text{nu}}$ in $\mu\text{m}^3$ | 2,9657 | 2,9309 | 3,0566 | 3,8402 | 3,8462 | 3,2246 | 3,7845 | 4,8302 | 4,1702 | 5,4659 | 6,8828 |
| rel. $V_{\text{nu}}$ in % | 0,1551 | 0,1348 | 0,1354 | 0,1567 | 0,1558 | 0,1302 | 0,1493 | 0,1745 | 0,1503 | 0,1547 | 0,1478 |

| RC3 | C5 | C6 | C8 | C1 | C4 | C2 | C11 | C12 | C9 | C3 | C10 | C7 |
| --- | --- | --- | --- | --- | --- | --- | --- | --- | --- | --- | --- | --- |
| abs. $V_{\text{cell}}$ in $\mu\text{m}^3$ | 10,1994 | 22,3389 | 24,2444 | 24,8660 | 26,3460 | 27,3644 | 30,3141 | 31,3991 | 32,1329 | 33,1629 | 35,6189 | 51,2403 |
| abs. $V_{\text{nu}}$ in $\mu\text{m}^3$ | 2,6216 | 3,7827 | 3,4616 | 3,8354 | 3,8246 | 3,8603 | 3,7898 | 5,1014 | 4,3098 | 4,8920 | 5,7396 | 8,6336 |
| rel. $V_{\text{nu}}$ in % | 25,70 | 16,93 | 14,28 | 15,42 | 14,52 | 14,11 | 12,50 | 16,25 | 13,41 | 14,75 | 16,11 | 16,85 |

| RC4 | C5 | C1 | C6 | C10 | C9 | C3 | C7 | C4 | C2 | C8 |
| --- | --- | --- | --- | --- | --- | --- | --- | --- | --- | --- |
| abs. $V_{\text{cell}}$ in $\mu\text{m}^3$ | 13,9838 | 18,0932 | 24,7512 | 25,7801 | 27,1277 | 30,9752 | 31,4192 | 31,4369 | 35,3603 | 36,4673 |
| abs. $V_{\text{nu}}$ in $\mu\text{m}^3$ | 4,2099 | 2,8101 | 3,2034 | 3,8192 | 3,8117 | 4,6823 | 4,8305 | 4,8008 | 5,4840 | 5,2734 |
| rel. $V_{\text{nu}}$ in % | 0,3011 | 0,1553 | 0,1294 | 0,1481 | 0,1405 | 0,1512 | 0,1537 | 0,1527 | 0,1551 | 0,1446 |

**Figure 3:**
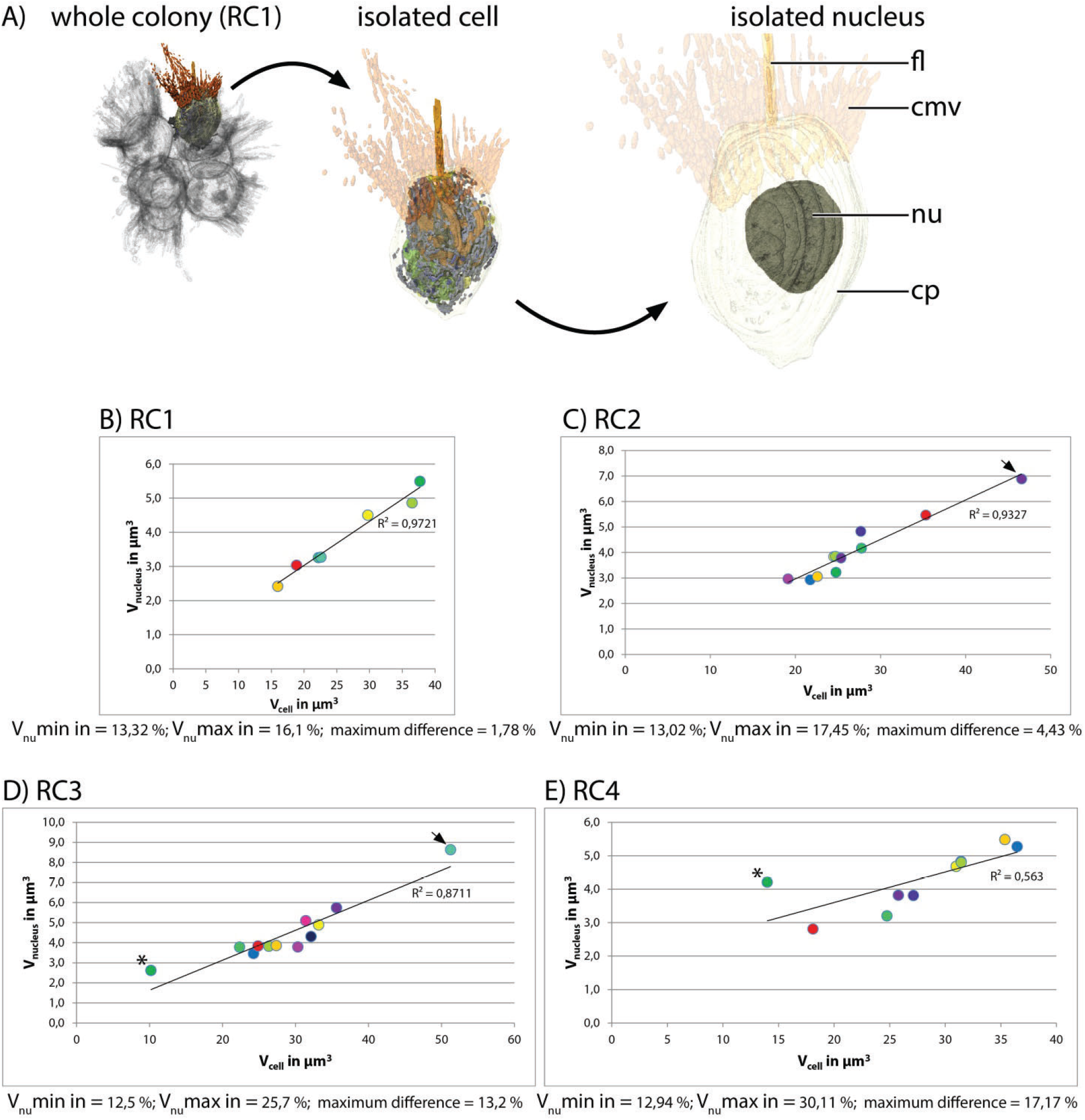
A, 3D-volume-renderings to illustrate the position and size of the nucleus in a colonial *S. rosetta* cell. B to E, plots of absolute nuclear volumes against the absolute cellular volume of cells from the four rosette colonies investigated in this study (RC1 to RC4). Cells are color coded according to Table 1. V_nu_max, maximal nuclear volume; V_nu_min, minimal nuclear volume.

In RC1 (Figure 3B), the mean volume of the nucleus relative to the total cell volume is 14,77%. C4 exhibits the lowest (13,32%) and C1 the highest relative nuclear volume (16,1%). Therefore, the maximum difference of the relative nuclear volumes between cells of RC1 is 2,78%. Comparing the absolute nuclear volumes C2, the smallest cell, exhibits the lowest (2,417 µm^3^) and C5, the largest cell, the highest absolute nuclear volume (5,495 µm^3^).

In RC2 (Figure 3C), the mean volume of the nucleus relative to the total cell volume is 14,95%. C7 exhibits the lowest (13,02%) and C9 the highest relative nuclear volume (17,45%). Therefore, the maximum difference of the relative nuclear volumes between cells of RC2 is 4,43%. Comparing the absolute nuclear volumes C8, the second smallest cell, exhibits the lowest (2,931 µm^3^) and C10, the largest cell, the highest absolute nuclear volume (6,883 µm^3^).

In RC3 (Figure 3D), the mean volume of the nucleus relative to the total cell volume is 15,9%. C11 exhibits the lowest (12,5%) and C5, the chili-shaped cell (Laundon et al. 2019), the highest relative nuclear volume (25,7%). Due to the exceptionally high nuclear volume of C5, the maximum difference of the relative nuclear volumes between cells of RC3 is 13,2%. Comparing the absolute nuclear volumes C5, the smallest cell, exhibits the lowest (2,622 µm^3^) and C7, the largest cell, the highest absolute nuclear volume (8,634 µm^3^).

In RC4 (Figure 3E), the mean volume of the nucleus relative to the total cell volume is 16,32%. C6 exhibits the lowest (12,94%) and C5, the carrot-shaped cell (Laundon et al. 2019), the highest relative nuclear volume (30,11%). As for RC3, due to the exceptionally high nuclear volume of C5, the maximum difference of the relative nuclear volumes between cells of RC3 is 17,17%. Comparing the absolute nuclear volumes C1, the second smallest cell, exhibits the lowest (2,81 µm^3^) and C2, the second largest cell, the highest absolute nuclear volume (5,484 µm^3^).

In summary, a relatively strong correlation between the nuclear volume and the total cell volume can be recognized (Figure 3B-E). In cells of RC1 (Figure 3B) and RC2 (Figure 3C) the correlation between nuclear volume and total cell volume is strongest. However, two “outlier cells”, C5 (Figure 3D; black asterisk) and C7 (Figure 3D; black arrow) are observed in RC3. In RC4, the correlation between nuclear and total cell volume is the lowest among the four colonies analyzed and an exceptional outlier cell, C5, is observed (Figure 3E; black asterisk). A plot of the relative mean, minimal and maximal nuclear volumes against colony size indicates a higher cell disparity in larger colonies (RC2, RC3 and RC4) compared to RC1 (maximum difference; Figure 3). However, intracolonial cell disparity seems not to increase in a stepwise manner. The minimal and mean relative nuclear volumes do not show a high variation between the colonies, most of the variation comes from the maximal relative nuclear volumes.

### Mitochondrial volume correlates with cell size in *S. rosetta* rosette colonies

Most mitochondria in single-cell and colonial *S. rosetta* are organized within a network, called the mitochondrial reticulum, surrounding the nucleus (Leadbeater 2015; Booth et al. 2018; Davis et al. 2019) (Figure 4A). Only the relative and absolute volume of mitochondria located in the cytoplasm of each cell of RC1 – RC4 were regarded as functional and reconstructed (Table 2). Mitochondria incorporated into food vacuoles were regarded as non-functional and were not reconstructed.

**Table 2:** Absolute and relative mitochondrial volumes of cells of RC1 to RC4. abs. V_cell_, absolute total cellular volume; abs. V_mt_, absolute mitochondrial volume; rel. V_mt_, relative mitochondrial volume.

| RC1 | C2 | C1 | C6 | C7 | C3 | C4 | C5 |
| --- | --- | --- | --- | --- | --- | --- | --- |
| abs. $V_{\text{cell}}$ in $\mu\text{m}^3$ | 15,9781 | 18,8541 | 22,1971 | 22,5861 | 29,7504 | 36,4994 | 37,7110 |
| abs. $V_{\text{mit}}$ in $\mu\text{m}^3$ | 1,0130 | 1,1019 | 1,1689 | 1,3063 | 1,7792 | 2,0567 | 2,1986 |
| rel. $V_{\text{mit}}$ in % | 6,34 | 5,84 | 5,27 | 5,78 | 5,98 | 5,63 | 5,83 |

| RC2 | C11 | C8 | C2 | C3 | C4 | C7 | C5 | C9 | C6 | C1 | C10 |
| --- | --- | --- | --- | --- | --- | --- | --- | --- | --- | --- | --- |
| abs. $V_{\text{cell}}$ in $\mu\text{m}^3$ | 19,1235 | 21,7361 | 22,5701 | 24,5022 | 24,6890 | 24,7695 | 25,3476 | 27,6794 | 27,7535 | 35,3241 | 46,5795 |
| abs. $V_{\text{mit}}$ in $\mu\text{m}^3$ | 1,2810 | 1,4604 | 1,1399 | 1,4480 | 1,5245 | 1,4907 | 1,3743 | 1,5678 | 1,7204 | 2,3561 | 2,8299 |
| rel. $V_{\text{mit}}$ in % | 6,70 | 6,72 | 5,05 | 5,91 | 6,17 | 6,02 | 5,42 | 5,66 | 6,20 | 6,67 | 6,08 |

| RC3 | C5 | C6 | C8 | C1 | C4 | C2 | C11 | C12 | C9 | C3 | C10 | C7 |
| --- | --- | --- | --- | --- | --- | --- | --- | --- | --- | --- | --- | --- |
| abs. $V_{\text{cell}}$ in $\mu\text{m}^3$ | 10,1994 | 22,3389 | 24,2444 | 24,8660 | 26,3460 | 27,3644 | 30,3141 | 31,3991 | 32,1329 | 33,1629 | 35,6189 | 51,2403 |
| abs. $V_{\text{mit}}$ in $\mu\text{m}^3$ | 0,4776 | 1,5074 | 1,3760 | 1,6103 | 1,7979 | 1,7390 | 2,0865 | 2,1444 | 1,9361 | 1,9053 | 2,2667 | 3,2181 |
| rel. $V_{\text{mit}}$ in % | 4,68 | 6,75 | 5,68 | 6,48 | 6,82 | 6,35 | 6,88 | 6,83 | 6,03 | 5,75 | 6,36 | 6,28 |

| RC4 | C5 | C1 | C6 | C10 | C9 | C3 | C7 | C4 | C2 | C8 |
| --- | --- | --- | --- | --- | --- | --- | --- | --- | --- | --- |
| abs. $V_{\text{cell}}$ in $\mu\text{m}^3$ | 13,9838 | 18,0932 | 24,7512 | 25,7801 | 27,1277 | 30,9752 | 31,4192 | 31,4369 | 35,3603 | 36,4673 |
| abs. $V_{\text{mit}}$ in $\mu\text{m}^3$ | 0,4290 | 1,0926 | 1,6646 | 1,5366 | 1,6422 | 2,0399 | 1,9526 | 2,0867 | 2,2740 | 2,3652 |
| rel. $V_{\text{mit}}$ in % | 3,07 | 6,04 | 6,73 | 5,96 | 6,05 | 6,59 | 6,21 | 6,64 | 6,43 | 6,49 |

**Figure 4:**
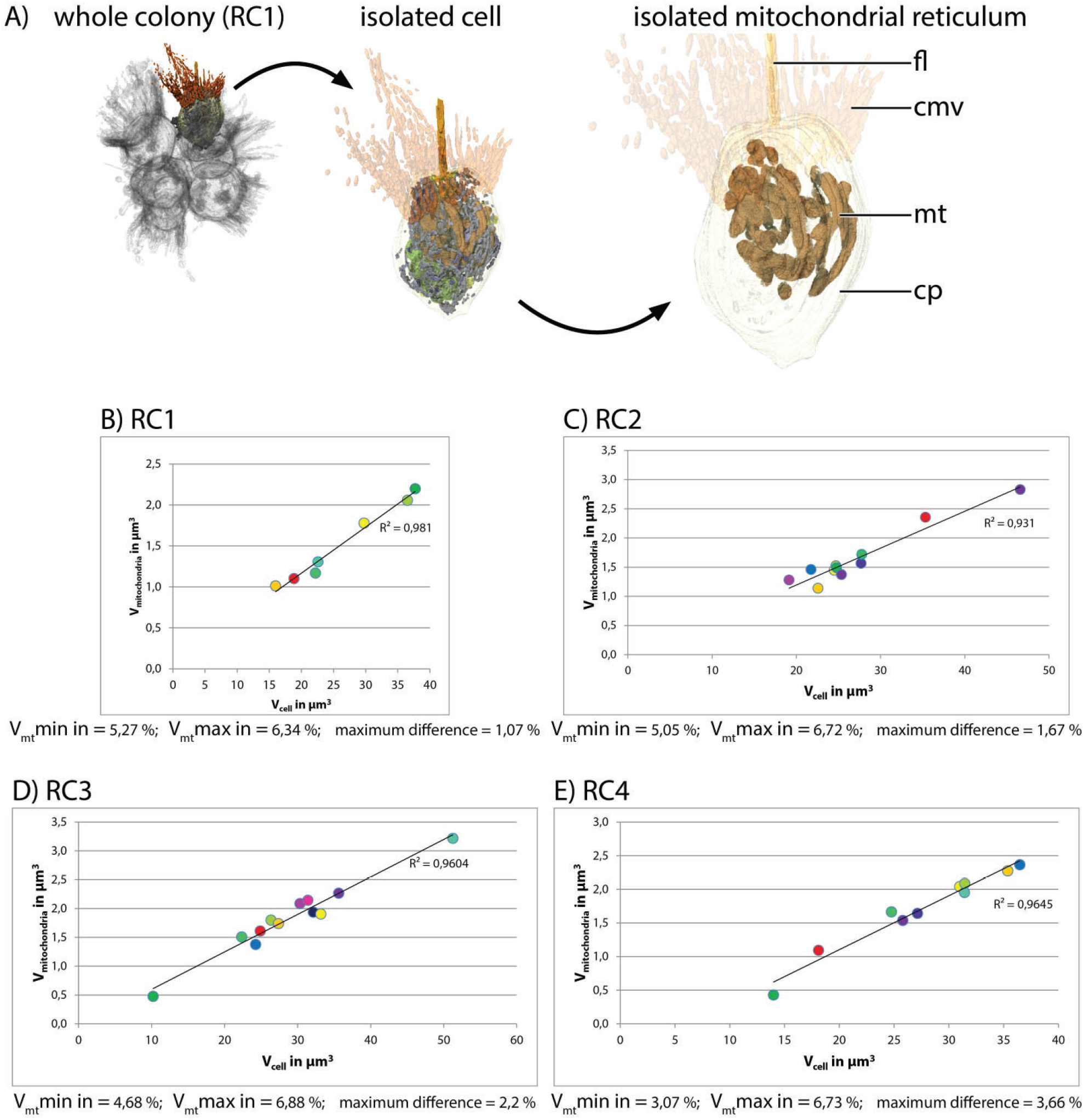
A, 3D-volume-renderings to illustrate the mitochondrial reticulum in a colonial *S. rosetta* cell. B to E, plots of absolute mitochondrial volumes against the absolute cellular volume of cells from the four rosette colonies investigated in this study (RC1 to RC4). Cells are color coded according to Table 2. V_mt_max, maximal mitochondrial volume; V_mt_min, minimal mitochondrial volume.

In RC1 (Figure 4B), the mean volume of the mitochondrial reticulum relative to the total cell volume is 5,81%. C6 exhibits the lowest (5,27%) and C2 the highest relative mitochondrial volume (6,34%). Therefore, the maximum difference of the relative mitochondrial volumes between cells of RC1 is 1,09%. Comparing the absolute mitochondrial volumes, C2, the smallest cell, exhibits the lowest (1,013 µm^3^) and C5, the largest cell, the highest absolute mitochondrial volume (2,197 µm^3^).

In RC2 (Figure 4C), the mean volume of the mitochondrial reticulum relative to the total cell volume is 6,05%. C2 exhibits the lowest (5,05%) and C8 the highest relative mitochondrial volume (6,72%). Therefore, the maximum difference of the relative mitochondrial volumes between cells of RC2 is 1,67%. Comparing the absolute mitochondrial volumes C2, the third smallest cell, exhibits the lowest (1,14 µm^3^) and C10, the largest cell, the highest absolute mitochondrial volume (2,83 µm^3^).

In RC3 (Figure 4D), the mean volume of the mitochondrial reticulum relative to the total cell volume is 6,24%. C5, the chili-shaped cell (Laundon et al. 2018), exhibits the lowest (4,68%) and C11 the highest relative mitochondrial volume (6,88%). Therefore, the maximum difference of the relative mitochondrial volumes between cells of RC3 is 2,2%. Comparing the absolute mitochondrial volumes, C5, the smallest cell, exhibits the lowest (0,478 µm^3^) and C7, the largest cell, the highest absolute mitochondrial volume (3,218 µm^3^).

In RC4 (Figure 4E), the mean volume of the mitochondrial reticulum relative to the total cell volume is 6,02 %. C5, the carrot-shaped cell (Laundon et al. 2018), exhibits the lowest (3,07 %) and C6 the highest relative mitochondrial volume (6,73 %). Therefore, the maximum difference of the relative mitochondrial volumes between cells of RC4 is 3,66 %. Comparing the absolute mitochondrial volumes, C5, the smallest cell, exhibits the lowest (0,429 µm^3^) and C8, the largest cell, the highest absolute mitochondrial volume (2,365 µm^3^).

In summary, our data indicate a strong correlation between the mitochondrial volume and the total cell volume in cells of each colony (Figure 4B-E). A plot of the relative mean, minimal and maximal mitochondrial volumes against the colony size (measured as cell number) indicates that the intracolonial cell disparity increases with the size of the colony (maximum difference; Figure 4). Most of this increase is due to a decrease of the relative minimal mitochondrial volume. The relative mean mitochondrial volume increases only slightly with colony size. The relative maximal mitochondrial volume is lowest in RC1 while almost similar in RC2, RC3 and RC4. We observed that in all colonies the majority of the mitochondria of a cell are organized as one large mitochondrial reticulum and only a few solitary mitochondria can be observed. However, an exact measurement of the number of mitochondria was not possible due to the section thickness of 150 nm (in RC2, RC3 and RC4). This thickness in combination with slight distortion artifacts from the sectioning process did not always allow reliable decisions as to whether one mitochondrium is continuous from one section to another or if it ends and another one begins in the following section.

### Food vacuole volume does not correlate with cell size in *S. rosetta* rosette colonies

In most cells food vacuoles are located in the basal half along the apical-basal axis of the cell. (Figure 5A). In the TEM sections analyzed, food vacuoles appear in different electron densities from high (dark grey) to relatively low (light grey). In between the two “extremes”, food vacuoles appear in different electron densities represented by different shades of grey. The electron density might represent different stages in the digestive cycle. To analyze the complete volume of food vacuoles within a cell we included all recognizable food vacuoles irrespective of their electron density (Table 3).

**Table 3:**
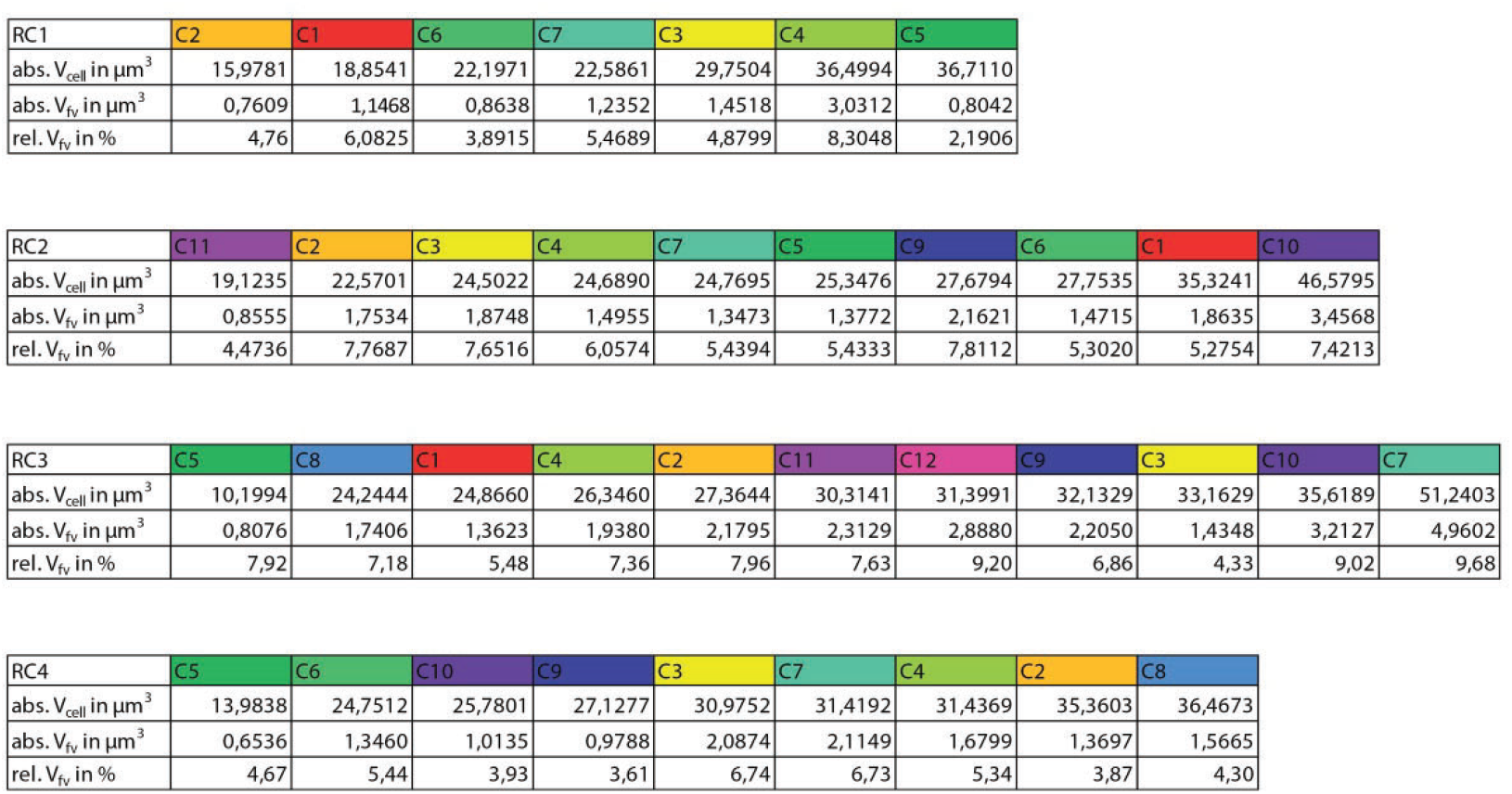
Absolute and relative food vacuole volumes of cells of RC1 to RC4. abs. V_cell_, absolute total cellular volume; abs. V_fv_, absolute food vacuole volume; rel. V_fv_, relative food vacuole volume.

**Figure 5:**
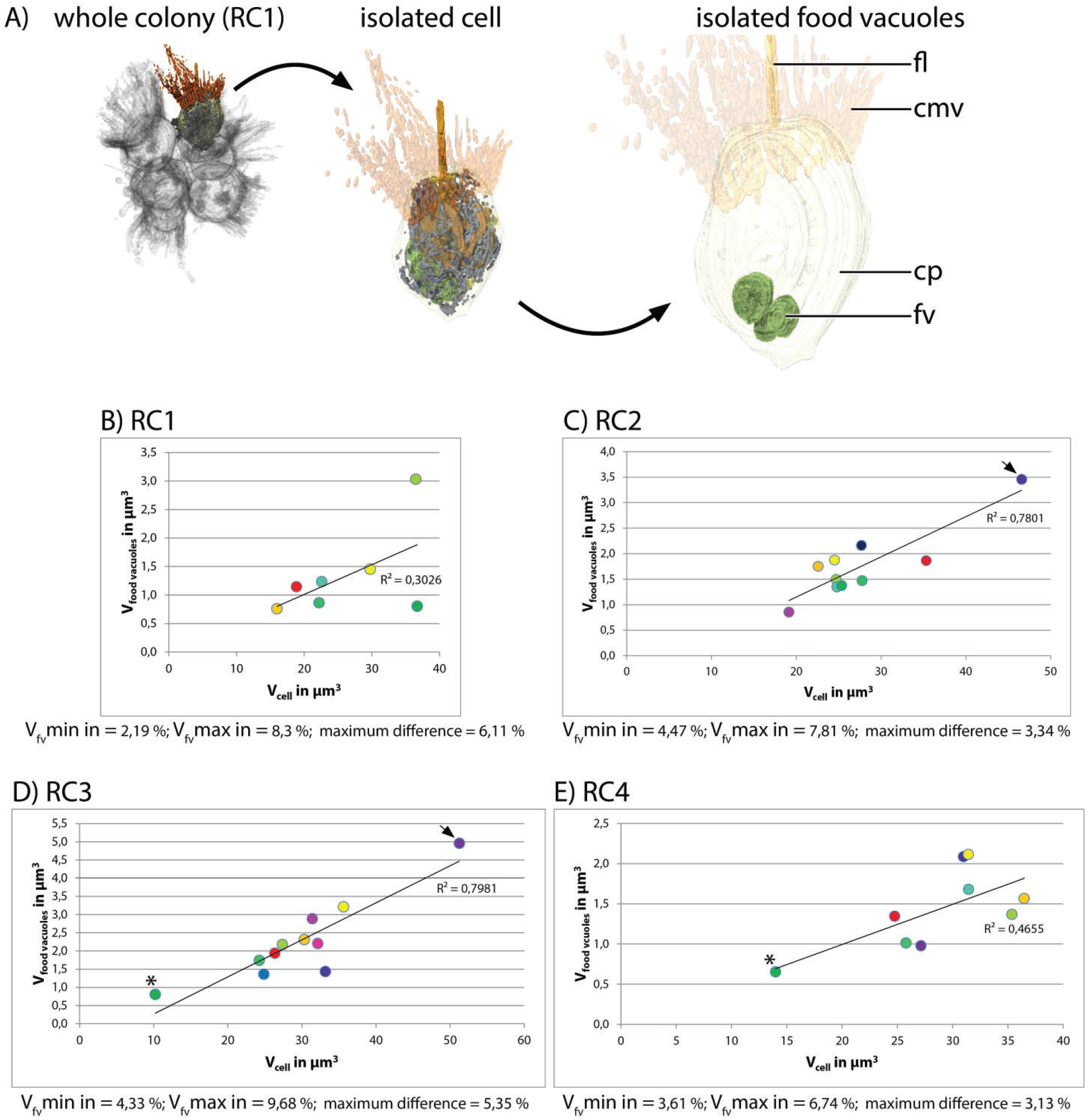
A, 3D-volume-renderings to illustrate the position and size of some food vacuoles in a colonial *S. rosetta* cell. B to E, plots of absolute food vacuole volumes against the absolute cellular volume of cells from the four rosette colonies investigated in this study (RC1 to RC4). Cells are color coded according to Table 3. V_fv_max, maximal food vacuole volume; V_fv_min, minimal food vacuole volume.

In RC1 (Figure 5B), the mean volume of food vacuoles relative to the total cell volume is 4,81%. C4, the largest cell, exhibits the lowest (2,19%) and C4, the second largest cell, the highest relative food vacuole volume (8,3%). Therefore, the maximum difference of the relative food vacuole volumes between cells of RC1 is 6,11%. Comparing the absolute food vacuole volumes, C2, the smallest cell, exhibits the lowest (0,761 µm^3^) and C4 the highest absolute food vacuole volume (3,031 µm^3^).

In RC2 (Figure 5C), the mean volume of food vacuoles relative to the total cell volume is 6,29%. C11, the smallest cell, exhibits the lowest (4,47%) and C3, the second smallest cell, the highest relative food vacuole volume (7,77%). Therefore, the maximum difference of the relative food vacuole volumes between cells of RC2 is 3,3%. Comparing the absolute food vacuole volumes C11, exhibits the lowest (0,856 µm^3^) and C10, the largest cell, the highest absolute food vacuole volume (3,457 µm^3^).

In RC3 (Figure 5D), the mean volume of food vacuoles relative to the total cell volume is 7,32%. C3, the third largest cell, exhibits the lowest (4,33%) and C7, the largest cell, the highest relative food vacuole volume (9,68%). Therefore, the maximum difference of the relative food vacuole volumes between cells of RC3 is 5,35%. Comparing the absolute food vacuole volumes C5, the smallest cell, exhibits the lowest (0,808 µm^3^) and C7 the highest absolute food vacuole volume (4,96 µm^3^).

In RC4 (Figure 5E), the mean volume of food vacuoles relative to the total cell volume is 4,93%. C9 exhibits the lowest (3,61%) and C3 the highest relative food vacuole volume (6,74%). Therefore, the maximum difference of the relative food vacuole volumes between cells of RC4 is 3,13%. Comparing the absolute food vacuole volumes, C5, the carrot-shaped cell (Laundon et al. 2018), exhibits the lowest (0,654 µm^3^) and C3 the highest absolute food vacuole volume (2,087 µm^3^).

In summary, our data indicate that there is only a weak correlation between food vacuole volume and the total cell volume in cells of each colony (Figure 5B-E). A plot of the relative mean, minimal and maximal food vacuole volumes against the colony size does not indicate that cell disparity changes with the size of the colony (maximum difference; Figure 5). However, our data indicate an increase of the minimal food vacuole volume with increasing cell size. An exact measurement of the number of food vacuoles was not possible due to the same limitations mentioned for the measurement of the mitochondrial number.

### Quantitative analysis of cell-cell contacts reveals plasma membrane contacts in colonial cells of *S. rosetta*

We found three different types of cell-cell contacts between cells in a colony (plasma membrane contacts, intercellular bridges and filo-/pseudopodial contacts; Figure 6). To our surprise, we discovered plasma membrane contacts between some cells of a colony (Figure 6A, B). These membrane contacts are found in all four colonies and range from relatively small areas (around 100 nm length on a section) up to areas of a length of more than 500 nm (length on a section). The presence of intercellular bridges (Figure 6A, C) and filo-/pseudopodial contacts (Figure 6A, D) has been described earlier (Dayel et al. 2011; Leadbeater 2015; Laundon et al. 2019).

**Figure 6:**
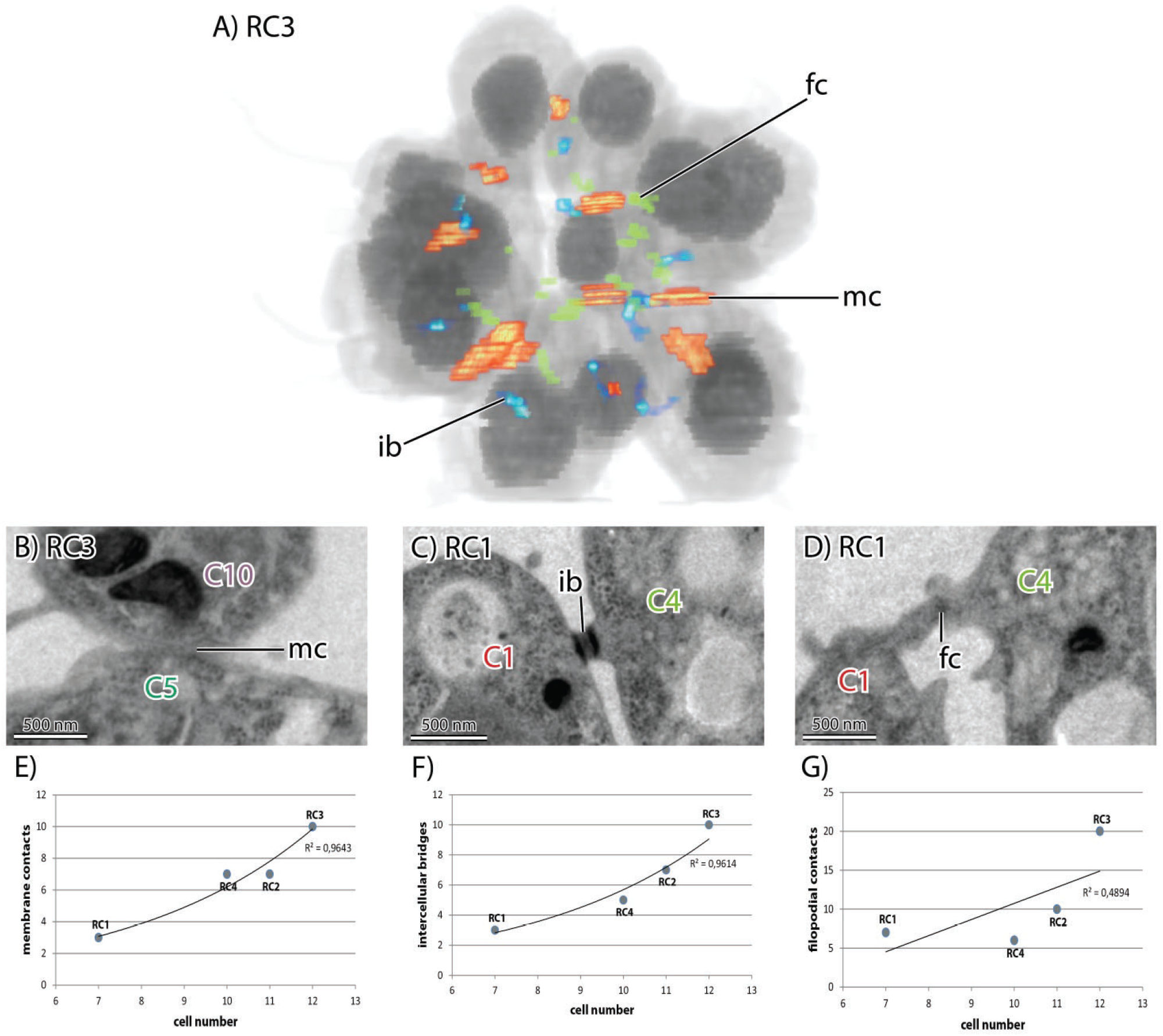
A, 3D-volume-rendering of *S. rosetta* rosette colony RC3 to illustrate the distribution of cell-cell contacts within a colony. B-D, TEM sections of different colonies highlight various types of cell-cell contacts in colonial *S. rosetta* cells. B, plasma membrane contact between C5 and C10 (RC3). C, intercellular bridge between C1 and C4 (RC1). D, filopodial contact between C1 and C4 (RC1). E-G, plots of the number of specific cell-cell contacts against the colony size (measured in cell number). ib, intercellular bridge; fc, filopodial contact; mc, membrane contact.

We quantified the number of the newly found plasma membrane contacts in the colonies used in this study. It seems that the number of plasma membrane contacts increases with the colony size (Fig. 6E). This is similar to the number of intercellular bridges (Figure 6F) and consistent with the results from Laundon et al. (2019). The number of filopodial/pseudopodial contacts between cells within the colonies seems not correlated with colony size (Figure 6G). A detailed summary of the types (intercellular bridges, membrane contact and filopodial contact) and number of connections of individual cell of each of the four investigated colonies is given in Table 4.

**Table 4:**
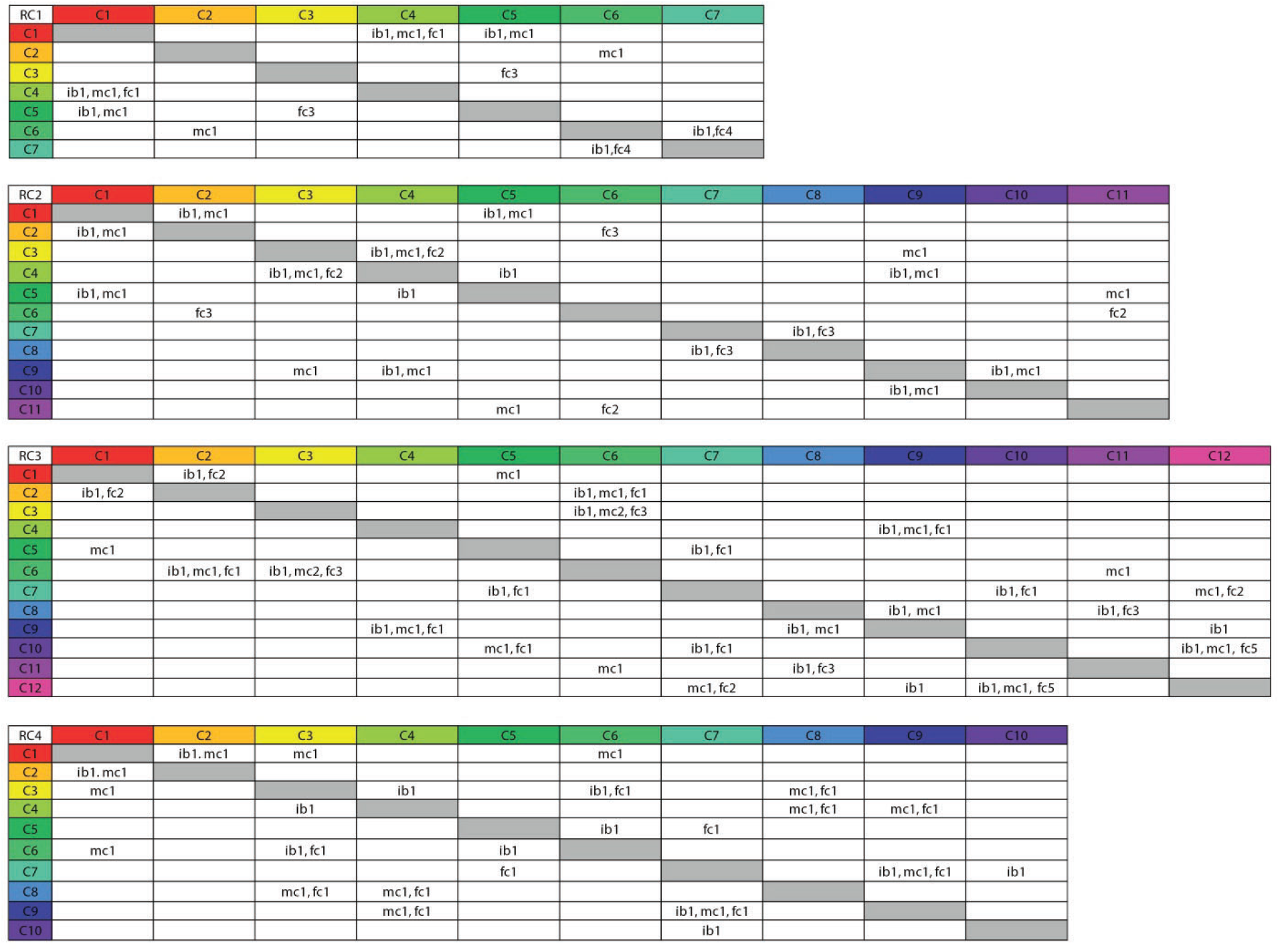
Types and of cell-cell contacts of cells of RC1 to RC4. Numbers behind the abbreviations indicate to total amount of the specific cell-cell contact. fc, filopodial-/pseudopodial contact; ib, intercellular bridge; mc, membrane contact.

## Discussion

In this study, we analyzed cell morphologies, volumes of cell bodies and volumes of some major organelles (nucleus, mitochondria and food vacuoles) of four *S. rosetta* rosette colonies (40 cells in total). The aims were: (1) To investigate whether cells in rosette colonies of *S. rosetta* are indeed “identical” or if they differ from each other. (2) In case they differ from each other, to what degree do they vary in terms of morphology, cell volume and organelle content? (3) To compare the intracolonial cell differences to the differences within a group of choanocytes of the homoscleromorph sponge *O. carmela*. The differences of cells within a colony are here described in a relative way using the term “cell disparity” (indicated by maximum volume differences in this study). Identical cells show no disparity at all, the maximum volume difference is zero. In contrast, cells with a high disparity exhibit a high maximum volume difference (Fig. 7).

**Figure 7:**
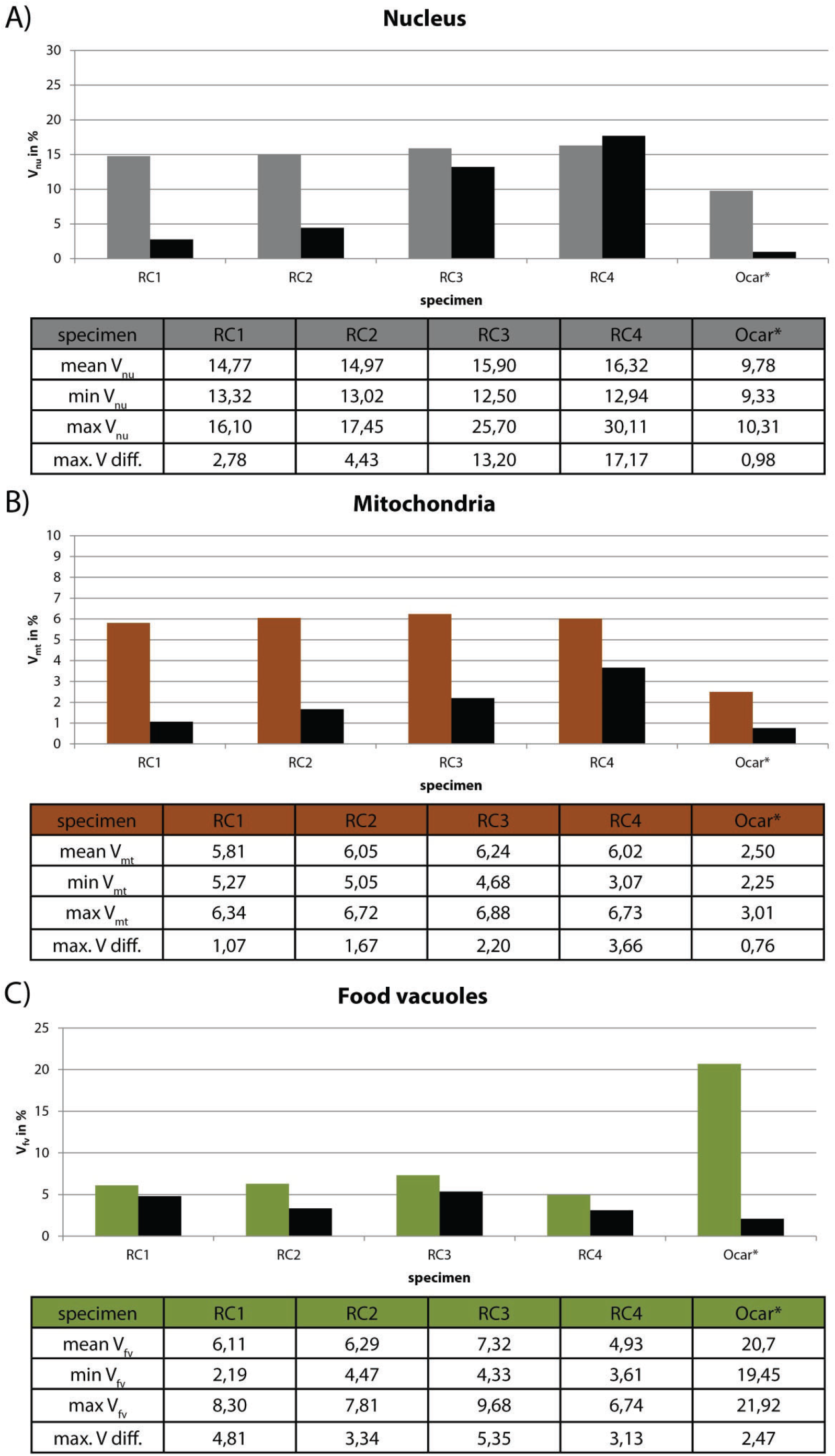
Plots of the relative volumes of different cell organelles of *S. rosetta* colonies (RC1 to RC4) and five choanocytes of *O. carmela* (Ocar). Values in the tables are given in % in relation to the total cellular volume. Asterisk, data taken from Laundon et al. (2019). A, nuclear volumes. B, mitochondrial volumes. C, food vacuole volumes. max. V diff., maximum volume difference; mean V, mean volume; Vmax, maximal volume; Vmin, minimal volume.

### Differences in nuclear volumes indicate a higher cell disparity within rosette colonies compared to sponge choanocytes

Size and form of the nucleus are often associated with cell division, differentiation, development and pathologies such as cancer. It is thought, that changes in nuclear size and form potentially impact chromatin organization, gene expression and other physiological processes of the cell (Jevtić et al. 2014). A study on myotube formation in human myoblasts has shown that a decrease in nuclear size is correlated with altered histone modifications, chromatin remodeling and gene silencing (Rozwadowska et al. 2013). The comparison of nuclear volumes of colonial cells (this study) with single cells of *S. rosetta* (Laundon et al. 2019) shows no difference between the two life history stages. This is in accordance with the almost identical transcriptomes of single cells and colonial cells of *S. rosetta* (Fairclough et al. 2013). However, comparison of the nuclear volume data of colonial *S. rosetta* cells with data from choanocytes of the sponge *O. carmel*a reveals interesting differences (Laundon et al. 2019). The relative nuclear volume of *O. carmela* choanocytes (9,78%, n = 5; Figure 7A) is around one third lower as the nuclear volume in cells of *S. rosetta* colonies (eg. RC1: 14,77%, n =7; Figure 7A) Additionally, the cell disparity among *O. carmela* choanocytes within the same choanocyte chamber of a sponge individual (maximum volume difference = 0,98%; Figure 7A) is much lower than among cells within one *S. rosetta* colony (eg. RC1: maximum volume difference = 2,78%; Figure 7A). These differences in the relative nuclear volume and cell disparity can be explained in two ways. The first explanation focuses on alterations of the nuclear volume due to cell division. For example, in the demosponge *Hymeniacidon sinapium*, choanocytes divide every 20 to 40 hours (Shore 1971). Cells of *S. rosetta* rosette colonies in contrast divide every 6 to 8 hours (Fairclough, Dayel, and King 2010). Due to the shorter cell cycle length of choanoflagellates compared to sponges it might be that the fixed *S. rosetta* colonies contained, by chance, more cells in the G2-phase of the cell cycle. G2-phase nuclei are often larger than nuclei during for example the G1-phase (Maeshima et al. 2011). The second explanation focuses on the specialized function of sponge choanocytes. Colonial choanoflagellate cells have been regarded as being more or less similar meaning that they show no real division of labor as present in Metazoa (Leadbeater 2015). Therefore choanoflagellate cells have to be “all-rounder” and constitutively express a variety of cellular modules such as for instance a ribosome biogenesis module, a flagellar module, a contractility module and a filopodia/microvilli module (Brunet and King 2017) to encounter all possible functional demands. Choanocytes in contrast, existing in a multicellular organism with other cell types, are specialized on only a few functions such as creating water current and food uptake (Simpson 2012; Mah et al. 2014; Dunn et al. 2015). Therefore, sponge choanocytes may not express a multitude of different cellular modules. Instead they might express only some modules in a cell type specific manner (Brunet and King 2017) (Fig.8). The expression of fewer cellular modules could be reflected by a decreased number of active genes and higher values of densely packed heterochromatin resulting in a smaller relative nuclear volume. Specialization could also explain the lower cell disparity in *O. carmela* choanocytes compared to colonial cells of *S. rosetta*. These arguments can be tested by investigating the chromatin architecture and euchromatin/heterochromatin ratios in “all-rounder” colonial cells of *S. rosetta* and specialized choanocytes of *O. carmela*.

**Figure 8:**
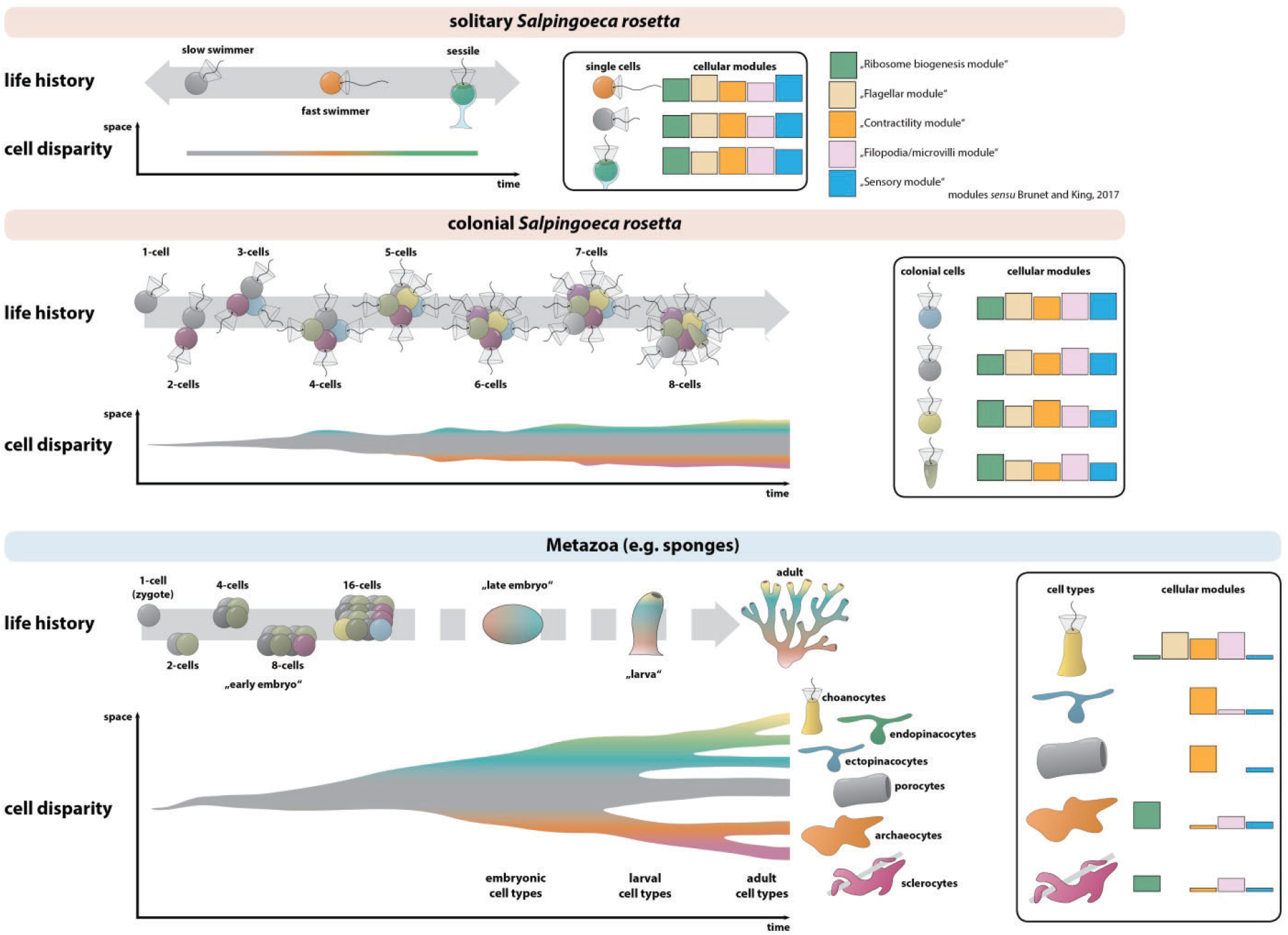
Hypothesis on spatial and temporal cell disparity in *S. rosetta* single cells, colonies and metazoans (e.g. sponges). Three cell types are described in solitary life history stages of *S. rosetta* (Dayel et al. 2011). Each of the solitary cell types might exhibit distinct expression levels of several constitutive cellular modules (*sensu* Brunet and King 2017) and cell disparity varies only in time but not in space. In colonial *S. rosetta*, cell disparity additionally varies in space. Upon increase of colony size the probability in cell disparity increases. In metazoans, cell number increases tremendously leading to a high degree of cell disparity. During development (and life history), cells differentiate into distinct cell types that express a specific set of cellular modules. This process decreases cell disparity between cells of the same cell types but increases cell disparity between cells belonging to different cell types.

### Differences of mitochondrial and food vacuole volumes indicate a higher cell disparity within rosette colonies compared to sponge choanocytes

Laundon et al. (2019) suggested, that the difference in mitochondrial number (single cells: 25,3% ± 5,8% *vs.* colonial cells: 4,3% ± 4,2%) and volume (single cells: 5,08% ± 1,14% *vs.* colonial cells: 6,63% ± 0,42 %) could be due to a higher demand on energy necessary for locomotion in single-cell *S. rosetta*. We confirm the results from Laundon et al. (2019) regarding the relative mean volume of the mitochondrial reticulum in colonial *S. rosetta* (RC1: 5,81%; RC2: 6,05%; RC3: 6,24%; RC4: 6,02%; Figure 7B). Our reconstructions of the mitochondrial reticulum of colonial cells also support the presence of a lower number of mitochondria in colonial cells (Laundon et al. 2019). However, it was not possible to determine the exact number of mitochondria in cells of RC2, RC3 and RC4 due to the thickness (150 nm) of the sections. It is known that mitochondrial fusion is stimulated by energy demand and stress while fission may generate new organelles and facilitates quality control (Youle and Van Der Bliek 2012). Limited mitochondrial fusion results in improper embryogenesis and is associated with some human diseases (Chen and Chan 2010). Therefore, fusion might act as a “defense mechanism” against cellular aging (Westermann 2002). Similar to our speculation on the variety of cell morphologies, the absence of extensive directed locomotion and the decreasing demand for high energy consumption in colonies might release cells from the constraint of having a high number of active, ATP-producing mitochondria. This may allow for a higher degree of mitochondrial fusion and increased longevity of mitochondrial function (Chen and Chan 2010). Laundon et al. (2019) reported a large difference in the mitochondrial volume of *S. rosetta* cells compared to choanocytes in *O. carmela*. We also confirm these results and show that the mean relative mitochondrial volume of colonial *S. rosetta* cells is around two times higher compared to sponge choanocytes (Figure 7B). As mentioned by Laundon et al. (2019) choanocytes as specialized cells without locomotory function might not need such high amounts of energy as free swimming *S. rosetta* cells with dual functions (food acquisition and locomotion) do. Additionally, we find that the cell disparity regarding mitochondrial volumes of cells within *S. rosetta* colonies (measured in maximum volume difference) is around 1,5 to 4,8 times higher compared to disparity within *O. carmela* choanocytes of the same individual (Figure 7B). The high cell disparity in choanoflagellate colonies could indicate early stages of “division of labor” (Bonner 2009; Brunet and King 2017). Cells of a colony are connected by intracellular bridges, pseudo-/filopodia (Dayel et al. 2011; Leadbeater 2015; Laundon et al. 2019) and membrane contacts (this study). Some of these structures could serve in exchange of metabolic compounds and could explain that certain cells within a colony can reduce the cellular volume devoted to mitochondria and increase the expression of other “cellular modules” (*sensu* Brunet and King 2017) while others increase the mitochondrial volume to cover the total energetic demands of the colony.

The high content of food vacuoles in sponge choanocytes (around 20,7% of total cell volume; Laundon et al. 2019) compared to the lower content in single-celled (around 9,22% of total cell volume; Laundon et al. 2019) and colonial (around 5,84% of total cell volume; this study) could also be explained by the principle of division of labor (Figure 7C). In single cell and colonial *S. rosetta* the dual function (food acquisition and locomotion) may demand for a trade-off between the cellular volume devoted to food vacuoles and other organelles, such as mitochondria, essential for flagellar function. While important for prey capture by producing a water current flagella of free-swimming single cell and colonial choanoflagellates are also involved in locomotion (Leadbeater 2015). The high levels of disparity of the relative food vacuole volume in the analyzed colonial cells of *S. rosetta* could be explained in several ways. First, these differences could arise simply from random differences of bacterial engulfment of individual cells. Another option is that some cells within the colony are more specialized on prey capture and distribute nutrients using cell-cell contacts. Such a transient specialization of groups of cells that facilitate locomotion and another group that can concentrate on prey capture might have been a key innovation in the metazoan stem lineage to escape the constraints imposed by the dual flagellar function. This “loss of constraints” hypothesis can be tested by analyzing the mitochondrial and food vacuole volumes of single cell *S. rosetta* under different prey concentrations and during different life history stages. Of special interest would be a comparison of the mitochondrial and food vacuole volumes between sessile, thecate *S. rosetta* and sponge choanocytes. If the “loss of constraints” hypothesis is correct, sessile *S. rosetta* should exhibit higher volumes of food vacuole and lower mitochondrial volumes than “slow and fast swimmer” cells and therefore be more similar to choanocytes.

### Plasma membrane contacts in *S. rosetta* rosette colonies

Cell-cell contacts and differential cell adhesion are key features during development and morphogenesis of metazoan embryos (Gilbert 2013). These contacts can be established in different ways utilizing intercellular bridges, filo-/pseudopodia and/or whole areas of the cell membrane. Intercellular bridges have been described in *S. rosetta* and several colony-forming choanoflagellates (Karpov and Coupe 1998; Dayel et al. 2011; Leadbeater 2015; Laundon et al. 2019). They have been hypothesized to function as channels for intercellular communication (Fairclough et al. 2013). It has been shown, that the number of intercellular bridges increases with the size of *S. rosetta* colonies (Laundon et al. 2018). In this study we report the presence of plasma membrane contacts between some cells of rosette colonies of *S. rosetta*. Our data show that the total number of intercellular plasma membrane contacts is comparable to the number of intercellular bridges and increases with colony size in a very similar pattern as observed for intercellular bridges (Figure 6A, B, D, E). Plasma membrane contact and cell adhesion in metazoans are mainly mediated by Cadherins (King et al. 2003; Cereijido et al. 2004; Halbleib and Nelson 2006). Twenty-three cadherins have been found in the strictly solitary choanoflagellate *Monosiga brevicollis.* Two of these cadherins localize in the microvillar collar and colocalize with the actin cytoskeleton (Abedin and King 2008). In *S. rosetta*, 29 proteins containing cadherin domains have been described (Nichols et al. 2012). However, the functions of these *S. rosetta* cadherins are still unknown (Fairclough et al. 2013). Some of the cadherins are differentially expressed during different stages of *S. rosetta* life history. Interestingly, two of these cadherins (PTSG_06458 and PTSG_06068) are upregulated in colonies compared to single cells (Fairclough et al. 2013). Further investigation of the spatial expression patterns of these two and other cadherins are crucial to clarify the properties and potential functions of intercellular membrane contacts in colonial choanoflagellates. In contrast to intercellular bridges and membrane contacts, the number of filo-/pseudopodial cell-cell contacts seems not tightly correlated with the size of a colony and might be a more variable and transient type of cell-cell contacts (Figure 6C, F). It remains to be examined if some more stable types of choanoflagellate cell-cell contacts (intercellular bridges, plasma membrane contacts) have homologous structures in metazoans and therefore might have been present in the last common ancestor of choanoflagellates and metazoans.

### Cell disparity, the evolution of metazoan cell differentiation and the last common ancestor of choanoflagellates and metazoans

Our data indicate, that (1) cells within a colony of *S. rosetta* are not identical but differ in terms of cell morphology, the relative volume ratios of several organelles (nucleus, mitochondria, food vacuoles) and their degree of interconnectedness within the colony. However, the degree to what they differ (2) varies. Colonies show a variety of cell morphologies from roundish over ovoid (along the AB-axis or horizontal to the AB-axis) to two extreme morphologies, the “carrot” (RC3; Laundon et al. 2019) and the “chili” cell (RC4; Laundon et al. 2019). All colonial cells show filo-, pseudo- and sometimes lobopodial membrane protrusion whose numbers seem to be independent from colony size (Figure 6). A comparison of the nuclear volumes indicates, that the cells in larger colonies (RC2: 11 cells; RC3: 12 cells; RC4: 10 cells) are more different (measured in maximal volume difference) compared to cells of the small colony (RC1: 7 cells) (Figure 7A). A similar trend can be observed regarding the mitochondrial volumes (Figure 7B). The volumes of food vacuoles, in contrast, seem more variable and show no colony size dependent pattern (Figure 7C).

Our study indicates that the cell disparity increases with the size of the choanoflagellate colony. The lion’s share of this cell disparity might be due to variations in metabolic processes (mitochondrial and food vacuole volumes) and cell cycle phases (nuclear and cell volumes), produced by different expression levels of constrictively expressed cellular modules (*sensu* Brunet and King 2017). The larger the colony the higher the statistical probability that cells exhibit different expression levels of cellular modules leading to higher degrees of cell disparity between them. Single cells for example only exhibit cell disparity in time (life history) but not in space because the same single cell can only have one specific identity at the time (Figure 8). A colony consisting of several cells can additionally exhibit cell disparity in space since different cells can have different identities. The more cells in a colony, the more cell identities can be present at the same time point leading to a higher possible cell disparity within the colony (Figure 8). However, a generalization of the idea that cell disparity increases with colony size is limited by the sample size investigated in this study. More *S. rosetta* colonies must be investigated in detail to test this idea. Another aim was (3) to compare our data to previously published data on nuclear, mitochondrial and food vacuole volumes in choanocytes of the homoscleromorph sponge *O. carmela* (Laundon et al. 2019). Our comparison supports the previous finding by Laundon et al. (2019) comparing three colonial *S. rosetta* cells to five choanocytes of *O. carmela*. *O. carmela* choanocytes exhibit smaller nuclear and mitochondrial but larger food vacuole volumes compared to cells of *S. rosetta* colonies (Laundon et al. 2019, this study; Figure 7). Additionally to these findings, we showed that the maximum volume difference of each of the three organelle volumes is much lower compared to colonial choanoflagellate cells (Figure 7). It seems that sponge choanocytes are not only more specialized on food acquisition (high volume of food vacuoles and lower mitochondrial and nuclear volumes) but also more similar to each other than individual cells in a colony of *S. rosetta*. Therefore, choanocytes exhibit lower spatial cell disparity compared to colonial *S. rosetta* cells (Figure 8). Is it possible to integrate this idea into an evolutionary context to explain the origin of metazoan cell types?

In contrast to the “Blastea/Gastrea” theory (Haeckel 1874, 1892), the Synzoospore hypothesis proposed that the origin of the Metazoa corresponds to the transition from temporal to spatial cell differentiation (Zakhvatkin 1949; Mikhailov et al. 2009). Zakhvatkin suggested that the last common ancestor of the Metazoa might have been an organism that already exhibited different cell types during different life history phases (temporal cell disparity and cell differentiation) as it can be seen in many protozoan taxa such as *S. rosetta* (Mikhailov et al. 2009; Dayel et al. 2011). During evolution, this organism acquired a benthic colonial or multicellular phase that was made up by cells of different cell types already present in the single cell stages of the life history of this organism. Mikhailov et al. (2009) suggested that it is unlikely that genetic programs of cell differentiation evolved *de novo* in this last common ancestor of the Metazoa. Instead, pre-existing mechanisms [cell differentiation programs] were used to integrate the different cell types that already occur during single cell life history phases of this organism. However, we think that it is more likely that new cell types evolved within a colony by co-option of cellular modules already present in cell types of single cell life history phases instead of including exactly the different cell types that already exist in single cell phases of the organism’s life history. This is indicated by the differences between solitary and colonial *S. rosetta* cells. On the basis of a detailed ultra-structural study, Laundon et al. (2019) suggested that colonial cells of *S. rosetta* might represent a distinct cell type instead of a conglomerate of identical “slow swimmer” cells. The “carrot” and “chili” cell may represent distinct cell types (Laundon et al. 2019), which is supported by our finding of high cell disparity in *S. rosetta* colonies.

Despite the controversy whether metazoans evolved from an ancestor exhibiting a “simple” or more complex life history, two main advantages have been proposed to drive positive selection for multicellularity in general. The first is an increase of size (Bonner 2009). Larger organisms/colonies might experience a lower predation pressure compared to smaller organisms/colonies (intercolonial competition) (Bonner 2009). However, after a certain size has been reached, cells within a colony might exhibit competition for space and nutrient availability (intracolonial competition). Therefore, selection might favor a better integration of cells by colonizing different “niches”, gradually becoming more different from each other (increasing spatial cell disparity) and eventually are recognized as different cell types. The result might have been a multicellular organism with different cell types that exhibit division of labor (Bonner 2009). Our investigation of cells within *S. rosetta* rosette colonies shows that the cells of a colony are not identical but differ from each other in terms of morphology and cell organelle volumes. The increased cell disparity in colonial life history phases might have been a pre-requisite for the colonization of different “intracolonial niches” by some cells leading to the evolution of division of labor by the differentiation into specialized cell types in the last common ancestor of the Metazoa.

## Acknowledgements

We thank Tarja Hoffmeyer, Ronja Goehde and Lennart Olsson for critical reading of the manuscript. This work was supported by the Sars Centre core budget.

---

**Supplementary Figure S1:** Scheme of the workflow and software types used in this study.

**Supplementary Figure S2:** 3D-surface-renderings of cells of a rosette colony of *S. rosetta* (RC1). Cells are not to scale. A, 3D-view of the whole colony from different angles. The color spectrum indicates the identity of the different cells. B-H, single views of cells of the colony. Cells are oriented along the apical (flagellar)-basal axis. The volume of the whole cell body is given beneath every cell.

**Supplementary Figure S3:** 3D-surface-renderings of cells of a rosette colony of *S. rosetta* (RC2). Cells are not to scale. A, 3D-view of the whole colony from different angles. The color spectrum indicates the identity of the different cells. B-L, single views of cells of the colony. Cells are oriented along the apical (flagellar)-basal axis. The volume of the whole cell body is given beneath every cell. C10 was not completely sectioned and half of the volume was added to estimate the total cellular volume indicated by the dotted line.

**Supplementary Figure S4:** 3D-surface-renderings of cells of a rosette colony of *S. rosetta* (RC3). Cells are not to scale. A, 3D-view of the whole colony from different angles. The color spectrum indicates the identity of the different cells. B-M, single views of cells of the colony. Cells are oriented along the apical (flagellar)-basal axis. The volume of the whole cell body is given beneath every cell.

**Supplementary Figure S5:** 3D-surface-renderings of cells of a rosette colony of *S. rosetta* (RC4). Cells are not to scale. A, 3D-view of the whole colony from different angles. The color spectrum indicates the identity of the different cells. B-K, single views of cells of the colony. Cells are oriented along the apical (flagellar)-basal axis. The volume of the whole cell body is given beneath every cell.

**Supplementary Figure S6:** 3D volume renderings of the nucleus, mitochondrial reticulum and food vacuoles of cells of a rosette colony of *S. rosetta* (RC1). Cells are not to scale. Cells are oriented along the apical (flagellar)-basal axis. The cell body is shown half transparent. The nucleus is colored in dark grey and the mitochondrial reticulum in brown. Food vacuoles with high electron density are colored in light green while food vacuoles with lower electron density are colored in dark grey.

**Supplementary Figure S7:** 3D volume renderings of the nucleus, mitochondrial reticulum and food vacuoles of cells of a rosette colony of *S. rosetta* (RC2). Cells are not to scale. Cells are oriented along the apical (flagellar)-basal axis. The cell body is shown half transparent. The nucleus is colored in dark grey and the mitochondrial reticulum in brown. Food vacuoles with high electron density are colored in light green while food vacuoles with lower electron density are colored in dark grey.

**Supplementary Figure S8:** 3D volume renderings of the nucleus, mitochondrial reticulum and food vacuoles of cells of a rosette colony of *S. rosetta* (RC3). Cells are not to scale. Cells are oriented along the apical (flagellar)-basal axis. The cell body is shown half transparent. The nucleus is colored in dark grey and the mitochondrial reticulum in brown. Food vacuoles with high electron density are colored in light green while food vacuoles with lower electron density are colored in dark grey.

**Supplementary Figure S9:** 3D volume renderings of the nucleus, mitochondrial reticulum and food vacuoles of cells of a rosette colony of *S. rosetta* (RC4). Cells are not to scale. Cells are oriented along the apical (flagellar)-basal axis. The cell body is shown half transparent. The nucleus is colored in dark grey and the mitochondrial reticulum in brown. Food vacuoles with high electron density are colored in light green while food vacuoles with lower electron density are colored in dark grey.

