## Supplementary figures and images for "Spatial cell disparity in the colonial choanoflagellate *Salpingoeca rosetta*"

### Supplemental Figures

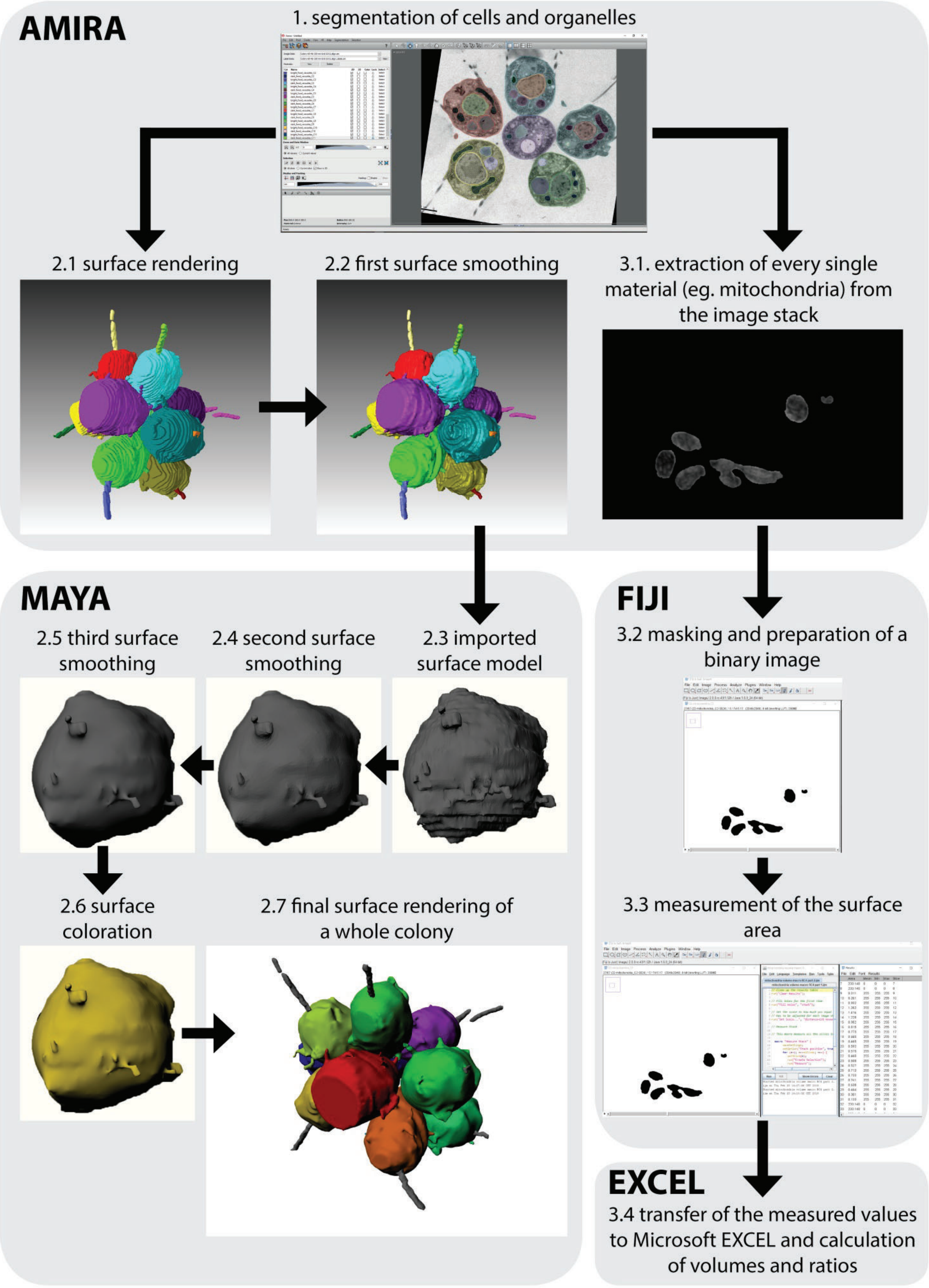

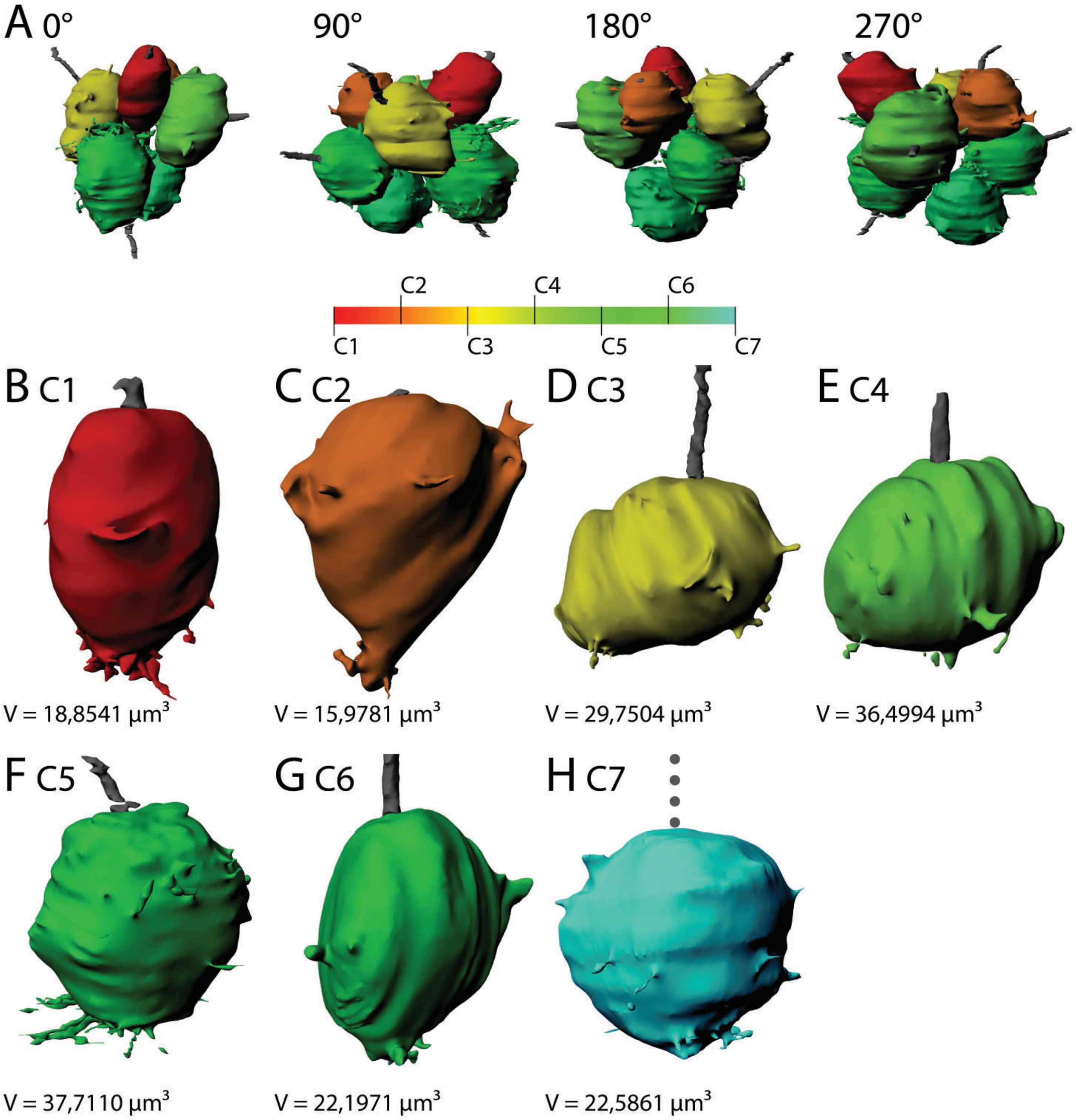

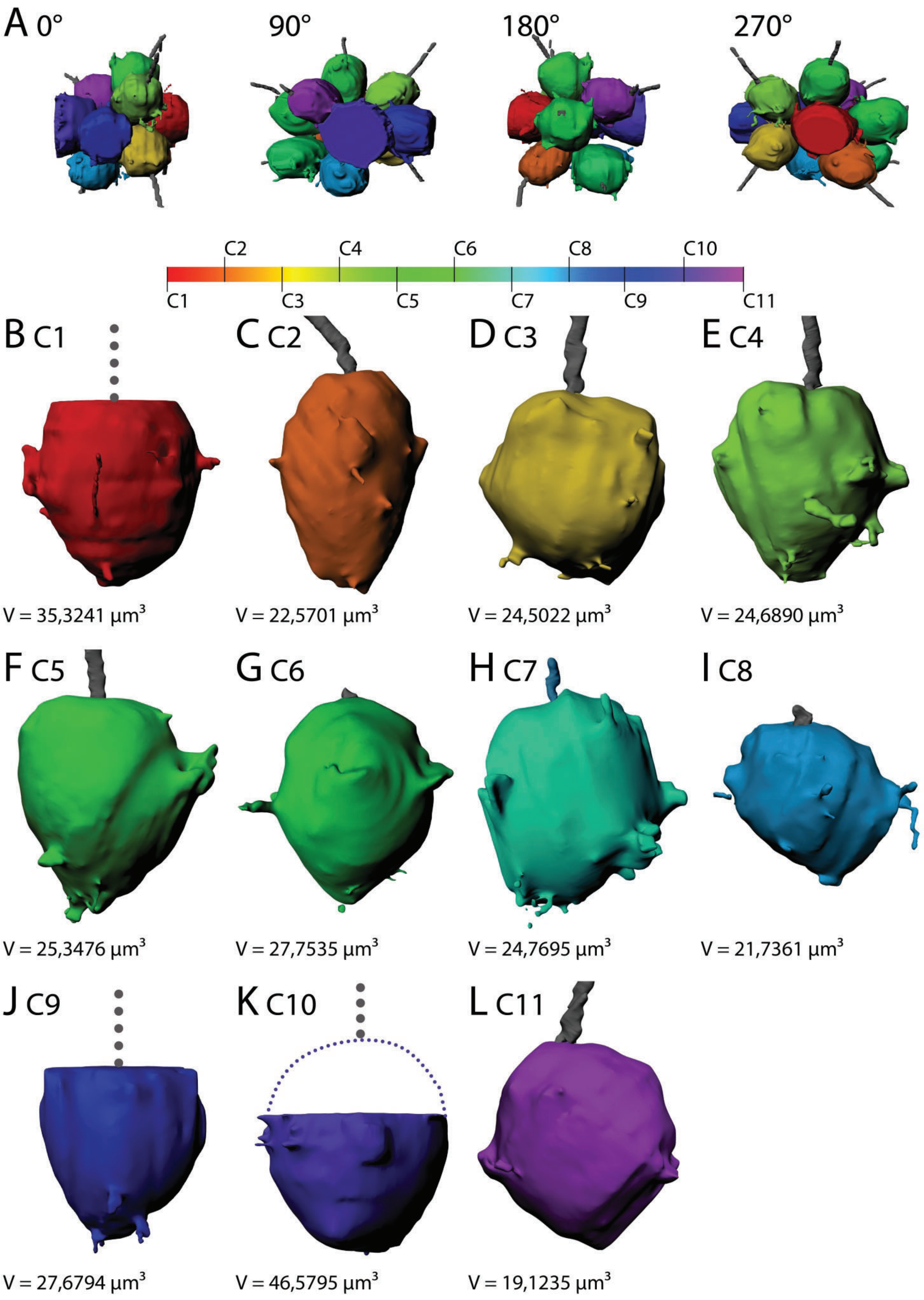

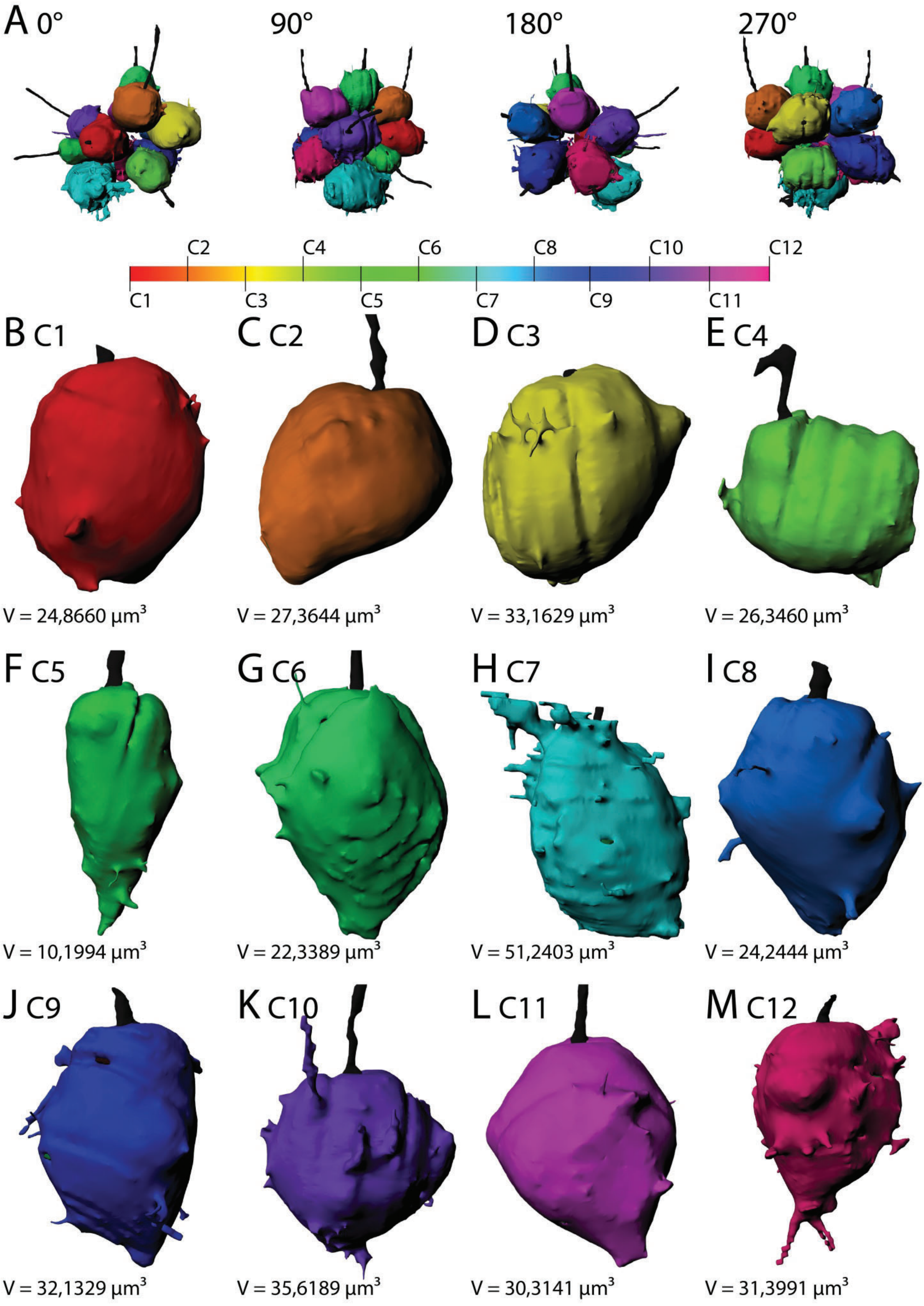

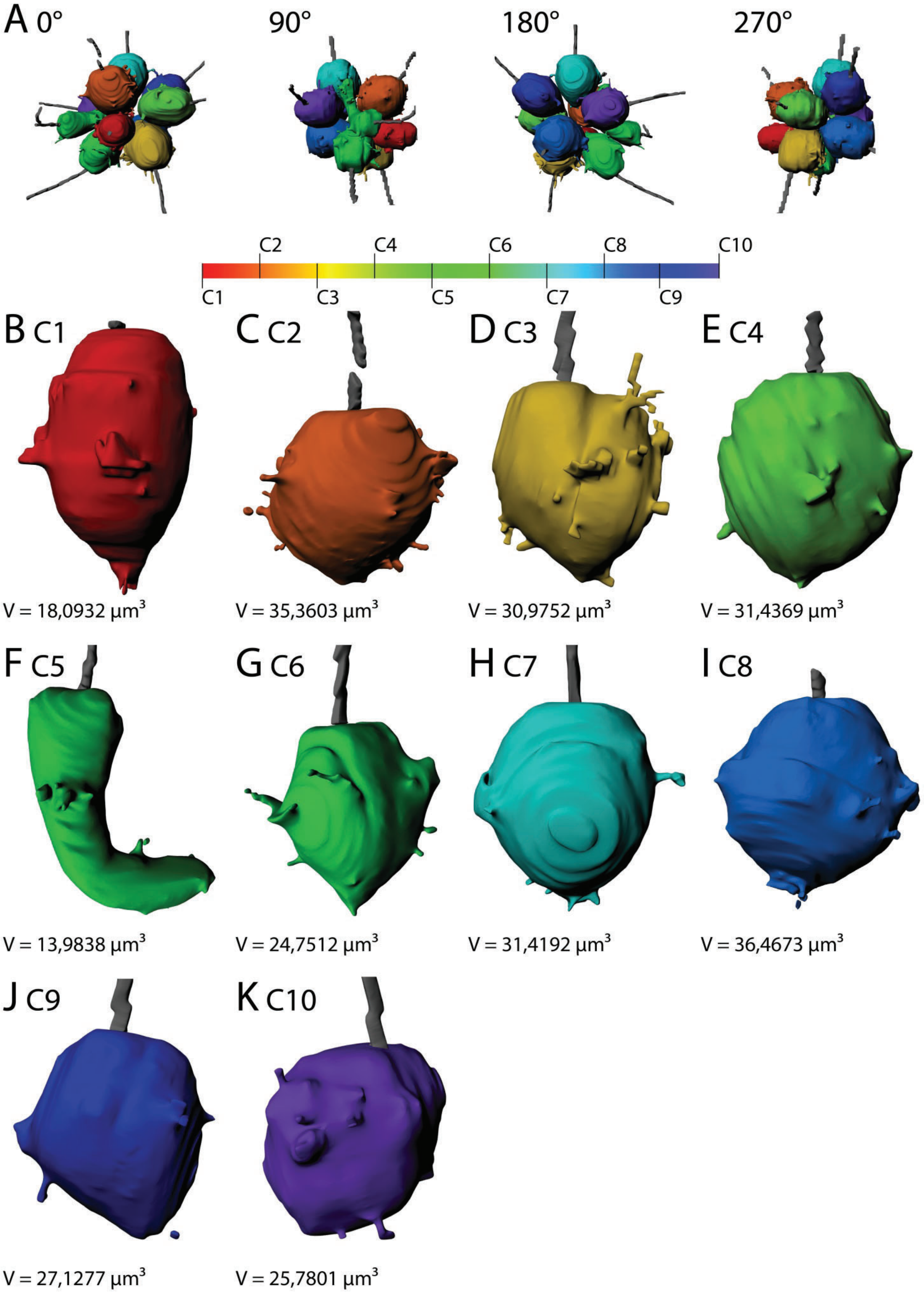

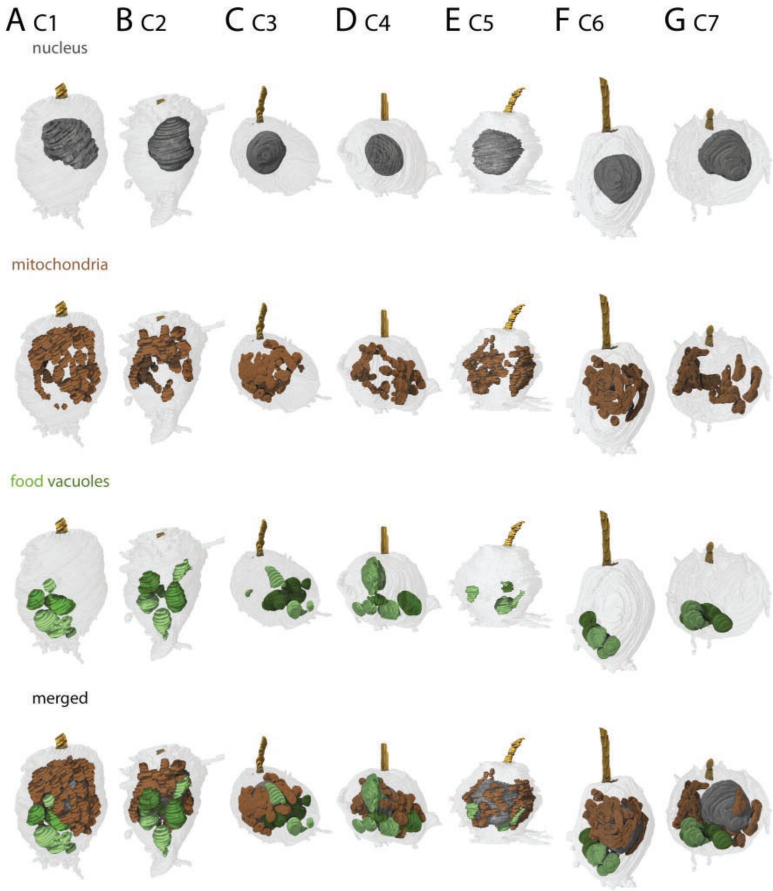

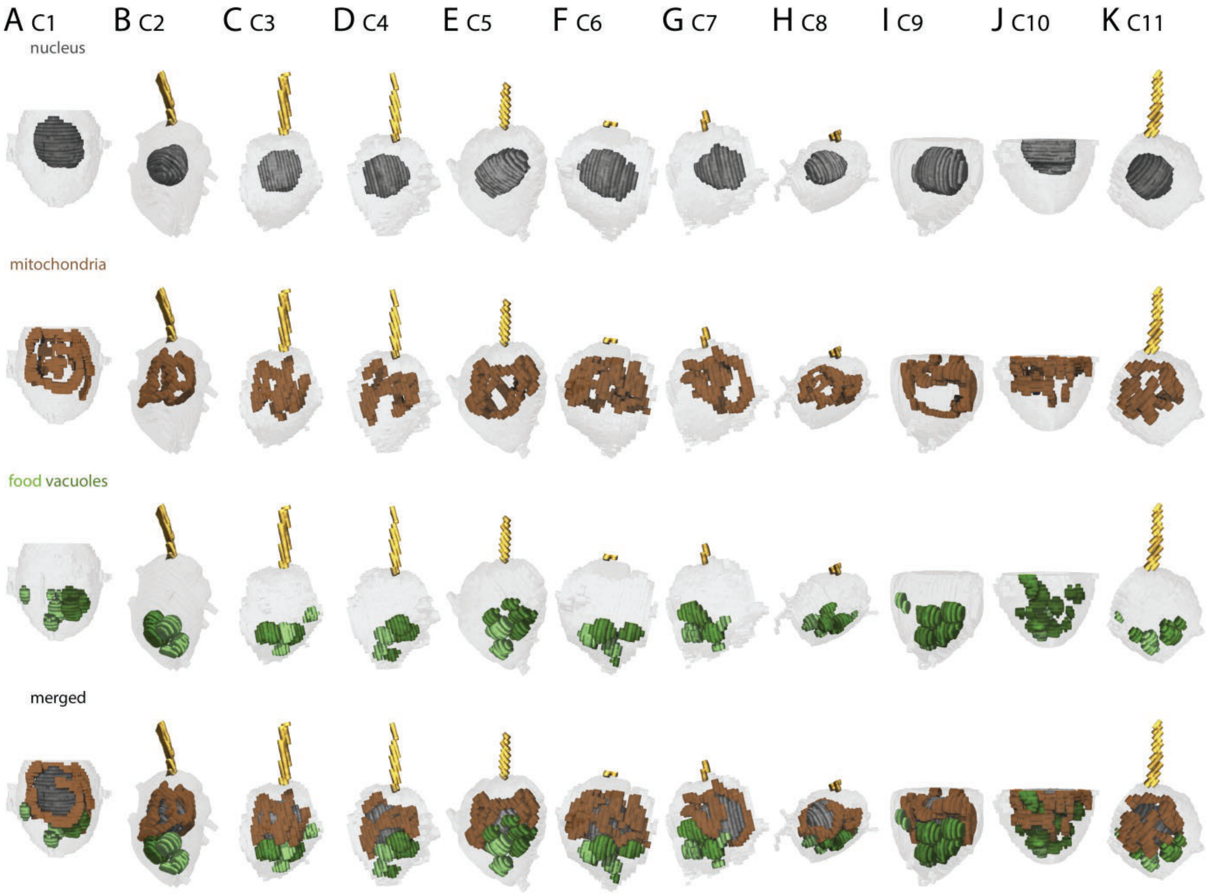

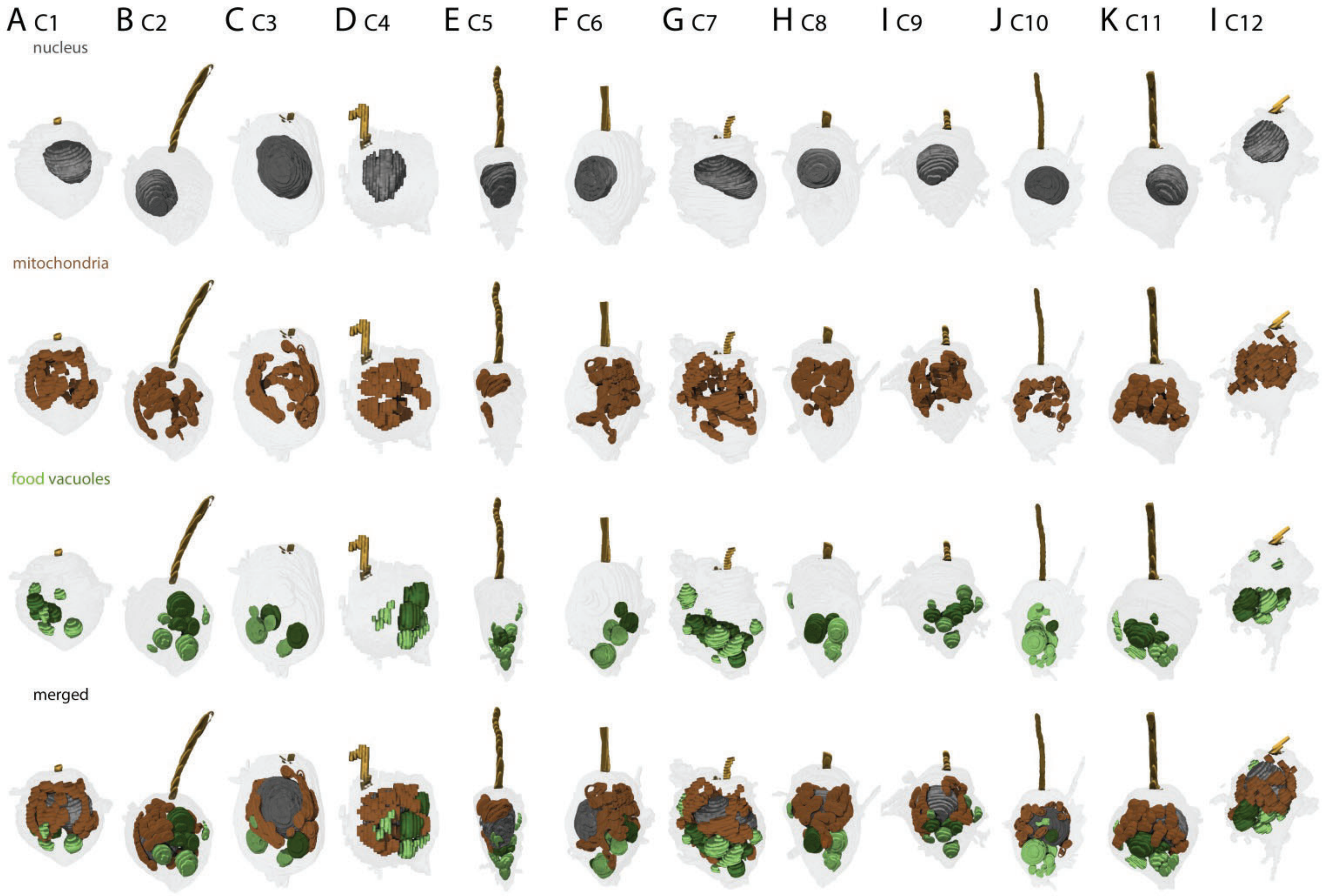

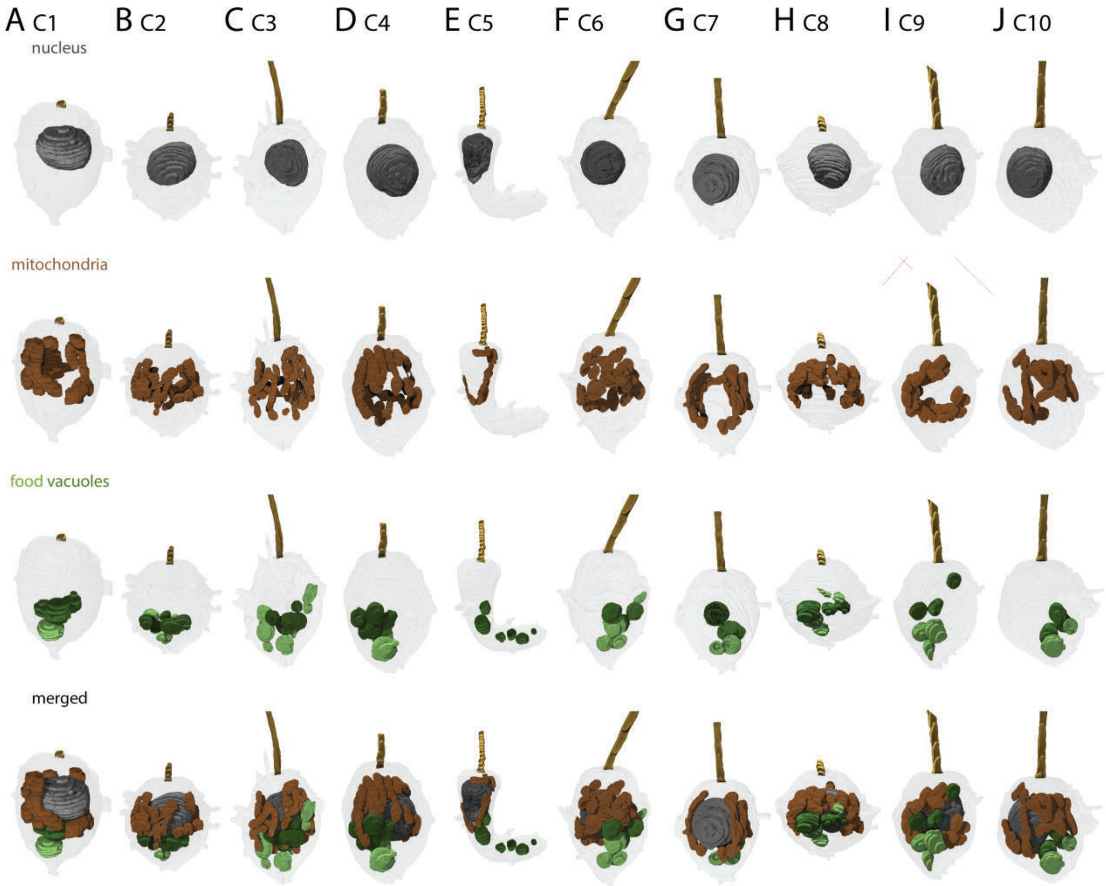
